# Two neuronal peptides encoded from a single transcript regulate mitochondrial complex III in *Drosophila*

**DOI:** 10.1101/2020.07.01.182485

**Authors:** Justin A. Bosch, Berrak Ugur, Israel Pichardo-Casas, Jorden Rabasco, Felipe Escobedo, Zhongyuan Zuo, Ben Brown, Susan Celniker, David A. Sinclair, Hugo Bellen, Norbert Perrimon

## Abstract

Naturally produced peptides (<100 amino acids) are important regulators of physiology, development, and metabolism. Recent studies have predicted that thousands of peptides may be translated from transcripts containing small open reading frames (smORFs). Here, we describe two peptides in *Drosophila* encoded by conserved smORFs, Sloth1 and Sloth2. These peptides are translated from the same bicistronic transcript and share sequence similarities, suggesting that they encode paralogs. Yet, Sloth1 and Sloth2 are not functionally redundant, and loss of either peptide causes animal lethality, reduced neuronal function, impaired mitochondrial function, and neurodegeneration. We provide evidence that Sloth1/2 are highly expressed in neurons, imported to mitochondria, and regulate mitochondrial complex III assembly. These results suggest that phenotypic analysis of smORF genes in *Drosophila* can provide a wealth of information on the biological functions of this poorly characterized class of genes.

## Introduction

Naturally produced peptides are regulators of metabolism, development, and physiology. Well-known examples include secreted peptides that act as hormones (Pearson *et al*. 1993), signaling ligands (Katsir *et al*. 2011), or neurotransmitters (Snyder and Innis 1979). This set of peptides are produced by cleavage of larger precursor proteins (Fricker 2005), peptides can also be directly translated from a transcript with a small open reading frame (smORF) (Couso and Patraquim 2017; Plaza *et al*. 2017; Hsu and Benfey 2018; Yeasmin *et al*. 2018). Due to their small size (<100 codons), smORFs have been understudied. For example, smORFs are underrepresented in genome annotations (Basrai *et al*. 1997), are theoretically a poor target for EMS mutagenesis, and are often ignored in proteomic screens. Consequently, there is growing interest in this class of protein-coding gene as a potentially rich source of novel bioactive peptides (Mudge *et al*. 2022).

A major obstacle in identifying smORFs that encode functionally important peptides is distinguishing them from the enormous number of smORFs present in the genome by chance (e.g. 260,000 in yeast) (Basrai *et al*. 1997). Many groups have identified and categorized smORFs with coding potential using signatures of evolutionary conservation, ribosomal profiling, and mass spectrometry (Saghatelian and Couso 2015; Couso and Patraquim 2017; Plaza *et al*. 2017).

Together, these approaches suggest there may be hundreds, possibly thousands, of unannotated smORF genes. However, these “omics” methods do not tell us which smORFs encode peptides with important biological functions.

Functional characterization of smORF genes in cell lines and model organisms has the potential to confidently identify novel peptides. Historically, unbiased genetic screens and gene cloning led to the fortuitous identification and characterization of smORF peptides (e.g. POLARIS (Casson *et al*. 2002), RpL41 (Suzuki *et al*. 1990), Nedd4 (Kumar *et al*. 1993), *Drosophila* pri/tal (Galindo *et al*. 2007)). More recently, candidate bioinformatically-predicted smORF-encoded peptides (aka SEPs) have been targeted for characterization (e.g., DWORF (Nelson *et al*. 2016), Elabela/toddler (Chng *et al*. 2013; Pauli *et al*. 2014), Myomixer (Bi *et al*. 2017), Myoregulin (Anderson *et al*. 2015), and Sarcolamban (Magny *et al*. 2013), and Hemotin (Pueyo *et al*. 2016)). Collectively, these studies have been invaluable for assigning biological functions to smORF peptides. Therefore, continued functional characterization is needed to tackle the enormous number of predicted smORF peptides.

Here, through an effort to systematically characterize human-conserved smORF genes in *Drosophila* (in preparation), we identified two previously unstudied smORF peptides CG32736-PB and CG42308-PA that we named Sloth1 and Sloth2 based on their mutant phenotypes. Remarkably, both peptides are translated from the same transcript and share amino acid sequence similarity, suggesting that they encode paralogs. Loss of function analysis revealed that each peptide is essential for viability, and mutant animals exhibit defective neuronal function and photoreceptor degeneration. These phenotypes can be explained by our finding that Sloth1 and Sloth2 localize to mitochondria and play an important role in complex III assembly. Finally, we propose that both peptides bind in a shared complex. These studies uncover two new components of the mitochondria and demonstrate how functional characterization of smORFs will lead to novel biological insights.

## Results

### *sloth1* and *sloth2* are translated from the same transcript and are likely distantly related paralogs

Current gene annotations for *sloth1* and *sloth2* (aka *CG32736* and *CG42308*, respectively) indicate that they are expressed from the same transcript (Flybase, Figure 1A), known as a bicistronic (or dicistronic) gene (Blumenthal 2004; Crosby *et al*. 2015; Karginov *et al*. 2017). For example, nearby transcription start sites (Figure 1A) are predicted to only generate a single transcript (Hoskins *et al*. 2011). In addition, a full-length transcript containing both smORFs is present in the cDNA clone RE60462 (GenBank Acc# AY113525), which was derived from an embryonic library (Stapleton *et al*. 2002), and we detected the full-length bicistronic transcript by RT-PCR amplification from total RNA from 3^rd^ instar larvae, adult flies, and S2R+ cells (Supplemental Figure 1). In addition, the encoded peptides Sloth1 and Sloth2 have subtle sequence similarity (27%), are similar in size (79aa and 61aa, respectively), and each contain a predicted single transmembrane domain (Figure 1B). While this type of gene structure is relatively rare in eukaryotes (Blumenthal 2004; Karginov *et al*. 2017), there are known cases in *Drosophila* of multicistronic transcripts encoding smORF paralogs – the *pri*/*tal* locus (Galindo *et al*. 2007) and the *Sarcolamban* locus (Magny *et al*. 2013). Furthermore, it is well known that paralogs are often found adjacent to each other in the genome due to tandem duplication (Taylor and Raes 2004). Therefore, we propose that *sloth1* and *sloth2* are paralogs translated from the same transcript.

**Figure 1:**
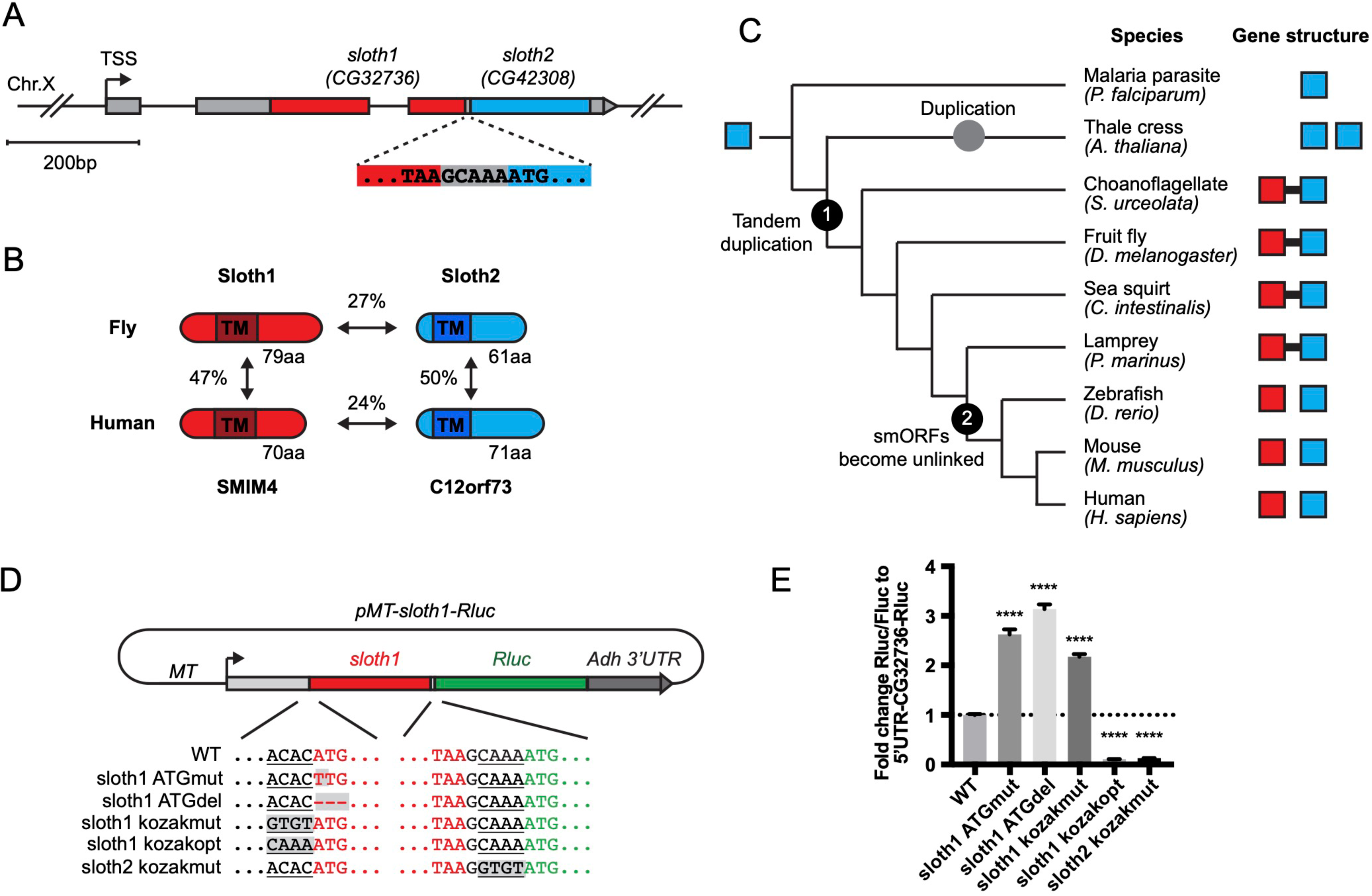
Bicistronic gene structure of the smORFs *sloth1* and *sloth2.* **A.** Bicistronic gene model for *sloth1* and *sloth2*. Zoom in shows intervening sequence (*GCAAA*) between *sloth1* stop codon and *sloth2* start codon. **B.** Comparison of protein structure, amino acid length size, and amino acid percent identity between *Drosophila* and Human orthologs. Shaded rectangle indicates predicted transmembrane (TM) domain. **C.** Phylogenetic tree of *sloth1* and *sloth2* orthologs in representative eukaryotic species. Linked gene structure (candidate bicistronic transcript or adjacent separate transcripts) is indicated by a black line connecting red and blue squares. **D.** Plasmid reporter structure of *pMT-sloth1- Rluc* and derivatives. Kozak sequences upstream of start codon are underlined. Mutations indicated with shaded grey box. pMT= Metallothionein promoter. RLuc = Renilla Luciferase. **E.** Quantification of RLuc luminescence/Firefly Luciferase, normalized to *pMT-sloth1-Rluc*, for each construct. Significance of mutant plasmid luminescence was calculated with a T-Test comparing to *pMT-sloth1- Rluc*. Error bars are mean with SEM. **** P≤0.0001. N=4 biological replicates.

Sloth1 and Sloth2 closely resemble their human orthologs (SMIM4 and C12orf73), based on sequence similarity, similar size, and presence of a transmembrane domain (Figure 1B). Like Sloth1 and Sloth2, SMIM4 and C12orf73 also have subtle amino acid sequence similarity to each other (Figure 1B). In addition, *sloth1* and *sloth2* are conserved in other eukaryotic species (Figure 1C). Remarkably, *sloth1* and *sloth2* orthologs in choanoflagelate, sea squirt, and lamprey exhibit a similar bicistronic gene architecture as *Drosophila* (Figure 1C, Supplemental File 1). In contrast, *sloth1* and *sloth2* orthologs in jawed vertebrates (e.g. mammals) are located on different chromosomes (e.g. human Chr.3 and Chr.12, respectively). Interestingly, we only found one ortholog similar to *sloth2* in the evolutionarily distant *Plasmodium*, and two orthologs similar to *sloth2* in *Arabidopsis*, which are located on different chromosomes (Figure 1C). Therefore, we hypothesize that the *sloth1* and *sloth2* ORFs duplicated from an ancient single common ancestor ORF and became unlinked in animals along the lineage to jawed vertebrates.

We next investigated *sloth1* and *sloth2* translation parameters and efficiency, since their ORFs are frameshifted relative to each other (Figure 1A) and they are not separated by an obvious internal ribosome entry site (IRES) (Van Der Kelen *et al*. 2009). Remarkably, only five nucleotides separate the stop codon of the upstream ORF (*sloth1*) and the start codon of the downstream ORF (*sloth2*) (Figure 1A). Therefore, *sloth1* should be translated first and inhibit translation of *sloth2*, similar to the functions of so-called upstream ORFs (uORFs) (Thompson 2012). However, *sloth1* has a non-optimal Kozak sequence 5’ to the start codon (*<u>ACAC</u>ATG*) and *sloth2* has an optimal Kozak (*<u>CAAA</u>ATG*) (Cavener 1987).

Therefore, scanning ribosomes may occasionally fail to initiate translation on *sloth1*, in which case they would continue scanning and initiate translation on *sloth2*, known as “leaky scanning” translation (Thompson 2012).

To test this translation model, we constructed an expression plasmid with the *Renilla Luciferase* (*RLuc*) reporter gene downstream of *sloth1* (*sloth1-RLuc*), while retaining non-coding elements of the original transcript (5’ UTR, Kozak sequences, 5bp intervening sequence) (Figure 1D). By transfecting this reporter plasmid into *Drosophila* S2R+ cells, along with a *Firefly Luciferase* (*FLuc*) control plasmid, we could monitor changes in translation of the downstream ORF by the ratio of RLuc/FLuc luminescence. Using derivatives of the reporter plasmid with Kozak or ATG mutations, we found that translation of the downstream ORF increased when translation of *sloth1* was impaired (Figure 1E). Reciprocally, translation of the downstream ORF was decreased when *sloth1* translation was enhanced with an optimal Kozak. These results suggest that *sloth1* inhibits translation of *sloth2*, and that balanced translation of both smORFs from the same transcript might be achieved by suboptimal translation of *sloth1*.

### *sloth1* and *sloth2* are essential in *Drosophila* with non-redundant function

To determine if *sloth1* and *sloth2* have important functions in *Drosophila*, we used in vivo loss of function genetic tools. We used RNA interference (RNAi) to knock down the *sloth1-sloth2* bicistronic transcript. Ubiquitous expression of an shRNA targeting the *sloth1* coding sequence (Figure 2A) lead to significant knockdown of the *sloth1-sloth2* transcript in 3^rd^ instar larvae (Figure 2B), as determined by two different primer pairs that bind to either the *sloth1* or *sloth2* coding sequence. Ubiquitous RNAi knockdown of *sloth1-sloth2* throughout development lead to reduced number of adult flies compared to a control (Figure 2C). This reduced viability was largely due to adult flies sticking in the food after they eclosed from their pupal cases (Figure 2D). Escaper knockdown flies were slow-moving and had 30% climbing ability compared to control flies (Figure 2E). RNAi knockdown flies also had short scutellar bristles (Figure 2F).

**Figure 2:**
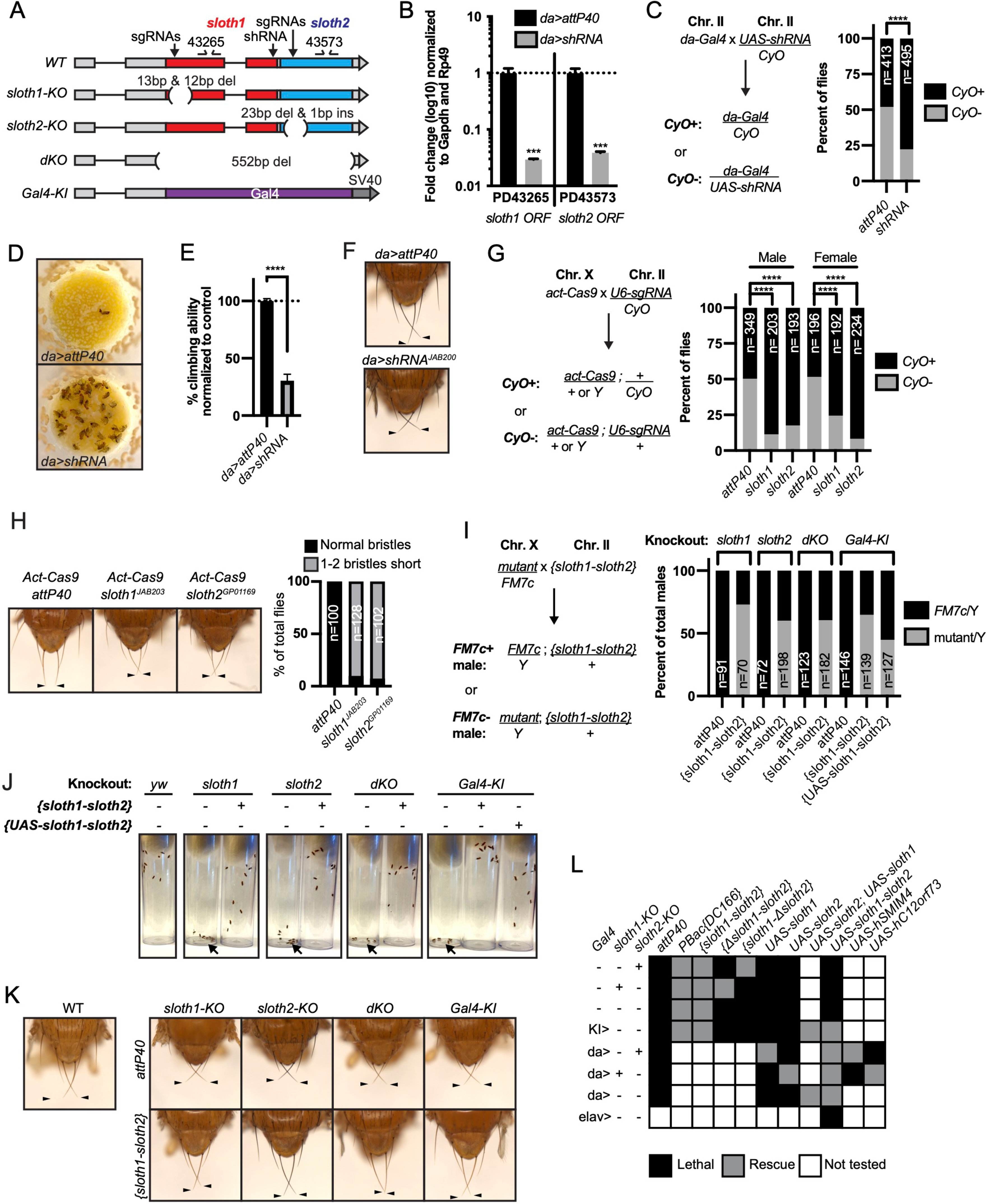
***sloth1* and *sloth2* loss of function analysis. A.** *sloth1-sloth2* transcript structure with shRNA and sgRNA target locations, primer binding sites, in/del locations, and knock-in Gal4 transgene. **B.** qPCR quantification of RNAi knockdown of the *sloth1*-*sloth2* transcript. Significance of fold change knockdown was calculated with a T-Test comparing to *da>attP40* for PD43265 and PD43573. Error bars show mean with SEM. P-values *** P≤0.001. N=6. **C.** Quantification of adult fly viability from *sloth1-sloth2* RNAi knockdown. Fly cross schematic (left) and graph (right) with percentage of progeny with or without the CyO balancer. Ratios of balancer to non-balancer were analyzed by Chi square test, **** P≤0.0001. Sample size (N) indicated on graph. **D.** Pictures of fly food vials, focused on the surface of the food. *da>shRNA* flies are frequently found stuck in the fly food. **E.** Quantification of adult fly climbing ability after *sloth1* and *sloth2* RNAi. Significance calculated with a T-test, **** P≤0.0001. Error bars show mean with SD. N=3 biological replicates. **F.** Stereo microscope images of adult fly thorax to visualize the scutellar bristles. RNAi knockdown by *da-Gal4* crossed with either *attP40* or *UAS-shRNA^JAB200^.* Arrowheads point to the two longest scutellar bristles. **G.** Quantification of adult fly viability from *sloth1-sloth2* somatic knockout. Fly cross schematic (left) and graph (right) with percentage of progeny with or without the CyO balancer. Ratios of balancer to non-balancer were analyzed by Chi square test, **** P≤0.0001. Sample size (N) indicated on graph. **H.** (Left) Stereo microscope images of adult fly thorax to visualize the scutellar bristles. Somatic knockout performed by crossing *Act-Cas9* to sgRNAs. (Right) Quantification of the frequency of adult flies with at least one short scutellar bristle after somatic KO of *sloth1* or *sloth2.* Sample sizes indicated on graph. Arrowheads point to the two longest scutellar bristles. **I.** Quantification of adult fly viability from *sloth1-sloth2* hemizygous knockout in males and rescue with a genomic transgene or *UAS-sloth1-sloth2* transgene. Fly cross schematic (left) and graph (right) with percentage of male progeny with or without the FM7c balancer. Sample size (N) indicated on graph. **J.** Still images from video of adult flies inside plastic vials. Images are 5 seconds after vials were tapped. Adult flies climb upward immediately after tapping. All flies are males. Each vial contains 10 flies, except dKO, which contains 5 flies. **K.** Stereo microscope images of adult male fly thorax to visualize the scutellar bristles. *attP40* is used as a negative control. Arrowheads point to the two longest scutellar bristles. **L.** Hemizygous mutant male genetic rescue experiments.

We confirmed our RNAi results using CRISPR/Cas9 to generate somatic knockout (KO) flies. By crossing flies ubiquitously expressing Cas9 (*Act-Cas9)* with flies expressing an sgRNA that targets the coding sequence of either *sloth1* or *sloth2* (Figure 2A, Supplemental Figure 2A), the resulting progeny will be mosaic for insertions and deletions (indels) that cause loss of function in somatic cells (Port *et al*. 2014; Xue *et al*. 2014). Both *sloth1* and *sloth2* somatic KO flies had significantly reduced viability compared to controls (Figure 2G). Furthermore, escaper adults had short scutellar bristles (Figure 2H) and frequently appeared sluggish. Importantly, similar phenotypes were observed when targeting either *sloth1* or *sloth2*.

Next, we further confirmed our loss of function results using CRISPR/Cas9 in the germ line to generate KO lines for *sloth1* and *sloth2*. These reagents are particularly important to test if *sloth1* and *sloth2* have redundant function by comparing the phenotypes of single and double null mutants. We generated four KO lines (Figure 2A, Supplemental Figure 2A-C): 1) a frameshift indel in *sloth1* (*sloth1*-KO), 2) a frameshift indel in *sloth2* (*sloth2*-KO), 3) a 552 bp deletion of the *sloth1* and *sloth2* reading frames (dKO), and 4) a knock-in of the reporter gene *Gal4* that removes *sloth1* and *sloth2* coding sequences (*Gal4-KI*). Since *sloth1* and *sloth2* are on the X-chromosome, we analyzed mutant hemizygous male flies. All four mutant lines were hemizygous lethal, which were rescued by a genomic transgene (Figure 2I,), ruling out off-target lethal mutations on the X-chromosome. Like RNAi and somatic KO results, rare mutant adult escaper flies had slower motor activity (Figure 2J) and short scutellar bristles (Figure 2K). Furthermore, the short scutellar bristle phenotype and slower motor activity could be rescued by a genomic transgene (Figure 2J, K).

The phenotypic similarity of single and double mutants suggests that *sloth1* and *sloth2* are not functionally redundant. However, since both ORFs are encoded on the same transcript, it is unclear if mutating one ORF will affect the other. For example, a premature stop codon can induce non-sense mediated decay of an entire transcript (Nickless *et al*. 2017). To address this possibility, we performed additional fly lethality rescue experiments. First, transheterozygous female flies (*sloth1-KO*/+, *sloth2-KO*/*+*) were viable and had normal scutellar bristles.

Second, we created single ORF versions of a genomic rescue transgene – *{Δsloth1-sloth2}* and {*sloth1*-Δ*sloth2*} (Supplemental Figure 2A). We found that *sloth1-KO* lethality could only be rescued by {*sloth1*-Δ*sloth2*}, and vice versa, *sloth2-KO* lethality could only rescued by {Δ*sloth1*-*sloth2*} (Figure 2L).

Furthermore, single ORF rescue transgenes were unable to rescue the lethality of *dKO* and *Gal4-KI* lines (Figure 2L). Third, we used the Gal4/UAS system (Brand and Perrimon 1993) to rescue mutant lethality with ubiquitously expressed cDNA transgenes. These results showed that single ORF KOs could only be rescued by expression of the same ORF (Figure 2L). Similar results were found by expressing cDNAs encoding the human orthologs (Figure 2L). In all, these results show that both *sloth1* and *sloth2* are essential, have similar loss of function phenotypes, are not functionally redundant with one another, and are likely to retain the same function as their human orthologs.

### Loss of *sloth1* and *sloth2* leads to defective neuronal function and degeneration

Since loss of *sloth1* and *sloth2* caused reduced adult mobility and climbing defects (Figure 2E, J), we speculated that the two peptides normally play an important role in the brain or muscle. To determine where *sloth1* and *sloth2* are expressed, we used the *Gal4-KI* line as an in vivo transcriptional reporter. *Gal4- KI* mobility defects and lethality could be rescued by expressing the entire bicistronic transcript (*UAS-sloth1-sloth2*) (Figure 2J, L), or coexpression of both smORFs as cDNA (*UAS-sloth1* and *UAS-sloth2)* (Figure 2L). Thus, the *Gal4-KI* line is likely an accurate reporter of *sloth1* and *sloth2* expression. By crossing *Gal4-KI* flies with a *UAS-GFP* fluorescent reporter, we observed strong GFP expression in larval (Figure 3A, B) and adult brains (Figure 3C). In addition, *Gal4- KI* is expressed in motor neurons at the larval neuromuscular junction (NMJ) (Figure 3D) and in larval brain cells that are positive for the neuronal marker Elav (Figure 3E).

**Figure 3.**
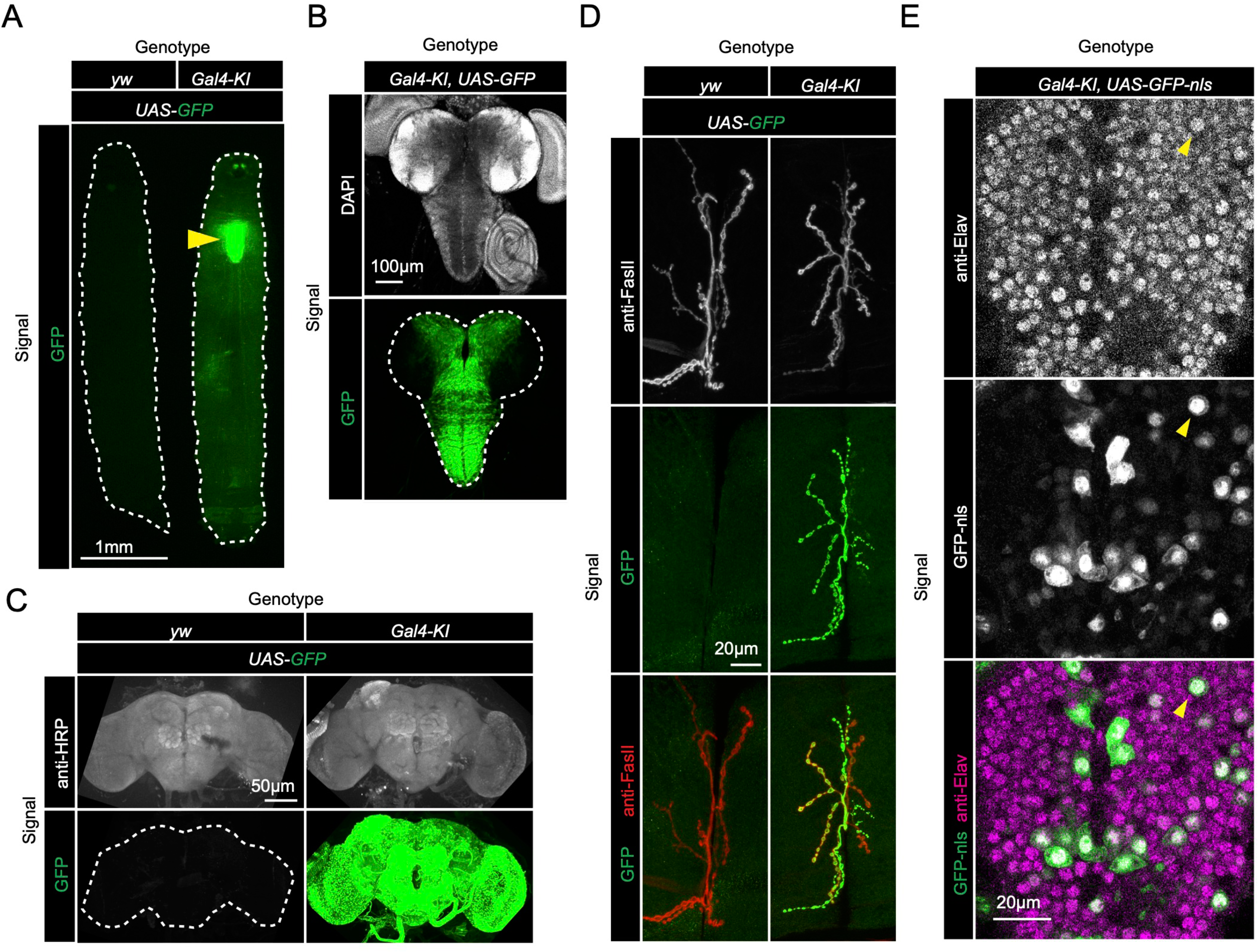
*sloth1*-*sloth2* are expressed in neurons. **A.** Fluorescent stereo microscope images of 3^rd^ instar larvae expressing GFP with indicated genotypes. **B.** Fluorescent compound microscope image of 3^rd^ instar larval brain expressing *UAS-GFP*. DAPI staining labels nuclei. **C.** Confocal microscopy of adult brain with indicated genotypes. Anti-HRP staining labels neurons. **D.** Confocal microscopy of the 3^rd^ instar larval NMJ at muscle 6/7 segment A2 expressing *UAS-GFP*. Anti-FasII staining labels the entire NMJ. **E.** Confocal microscopy of the 3^rd^ instar larval ventral nerve cord (VNC) expressing *Gal4-KI*, *UAS-GFP-nls*. GFP-nls is localized to nuclei. Anti-Elav stains nuclei of neurons. Arrow indicates example nuclei that expresses UAS-GFP and is positive for Elav.

We then tested if *sloth1* and *sloth2* were important for neuronal function by measuring neuronal electrical activity in *dKO* animals. Electrical recordings taken from the larval NMJ showed that *dKO* motor neurons have normal excitatory junction potential (EJP) under resting conditions at 0.75 mM Ca ^2+^ (Supplemental Figure 3). However, under high frequency stimulation (10hz), *dKO* NMJs could not sustain a proper response (Figure 4A), indicating that there is a defect in maintaining synaptic vesicle pools. Importantly, this phenotype is rescued by a genomic transgene. To test if a similar defect is present in the adults, we assessed phototransduction and synaptic transmission in photoreceptors via electroretinogram (ERG) recordings (Wu and Wong 1977; Hardie and Raghu 2001). ERGs recorded from young (1-3 days old) *dKO* photoreceptors showed an amplitude similar to that of genomic rescue animals (Figure 4B). However, upon repetitive light stimulation, ERG amplitudes were significantly reduced (Figure 4B), suggesting a gradual loss of depolarization. Similar results were observed when young flies were raised in 24hr dark (Figure 4C). Moreover, ERG traces also showed a progressive loss of “on” and “off” transients (Figure 4B, C), which is indicative of decreased synaptic communication between the photoreceptor and the postsynaptic neurons. ERG phenotypes are rescued by a full-length genomic rescue transgene, but not by single ORF rescue transgenes (Figure 4B, C). To test if loss of both *sloth1* and *sloth2* lead to neurodegeneration, we aged the animals for 4-weeks in 12hr light/dark cycle or constant darkness and recorded ERGs. Similar to young animals, aged animals raised in light/dark conditions also displayed a reduction in ERG amplitude upon repetitive stimulation (Figure 4E). These results indicate that both *sloth1* and *sloth2* are required for sustained neuronal firing in larval motor neurons and adult photoreceptors. Interestingly, similar mutant phenotypes in the NMJ and photoreceptors are known to be due to defects in ATP production (Verstreken *et al*. 2005; Sandoval *et al*. 2014; Jaiswal *et al*. 2015).

**Figure 4.**
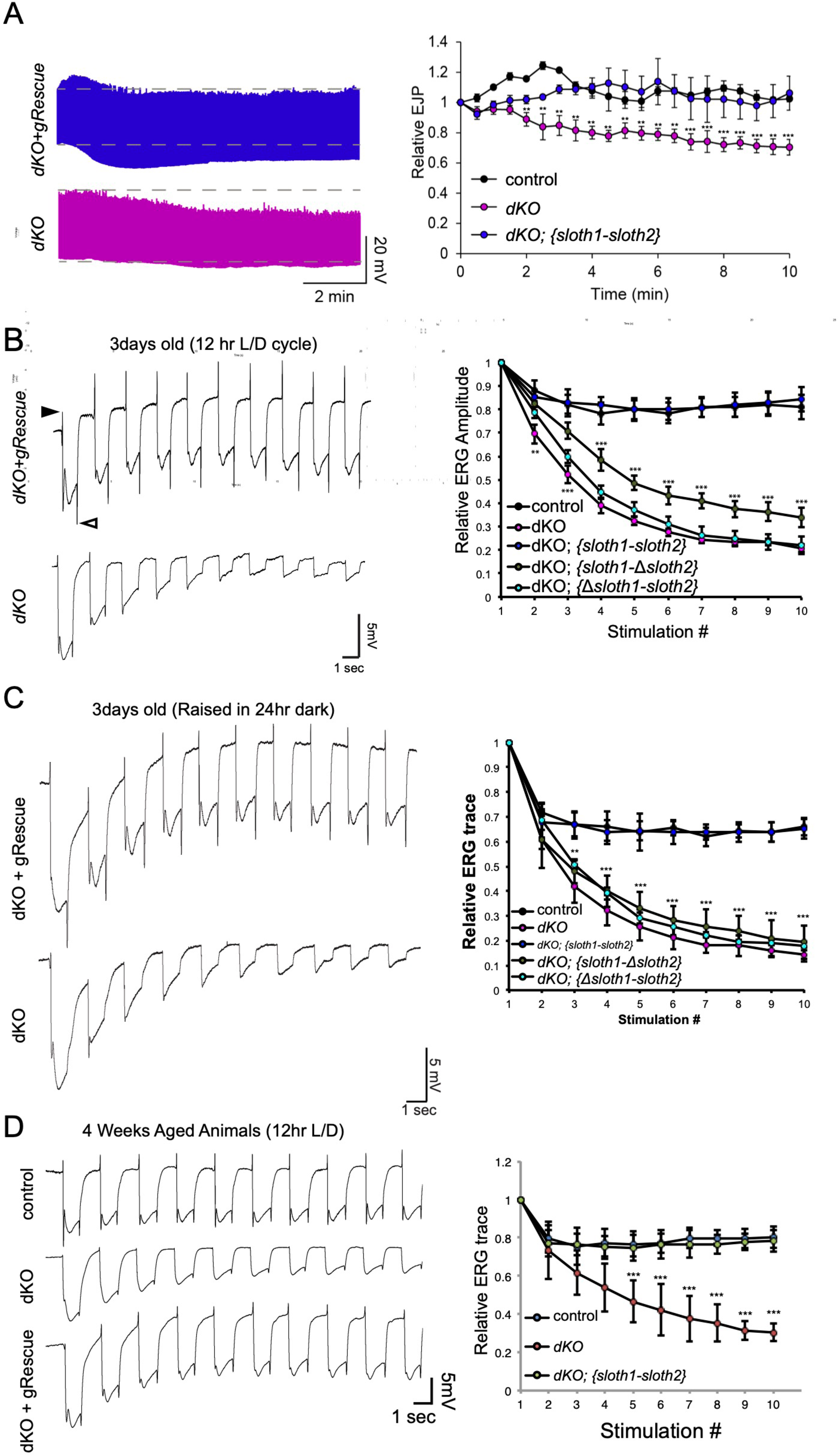
*sloth1*-*sloth2* are important for neuronal function. **A.** Traces of electrical recordings from 3^rd^ instar larval NMJ in control, *dKO*, and *dKO+genomic rescue* animals over 10 minutes under high frequency stimulation (10 Hz). Graph on right is a quantification of the relative excitatory junction potential (EJP) for indicated genotypes. Error bars show mean with SD. N ≥ 5 larvae per genotype. Significance for each genotype was calculated with a T-Test comparing to control flies. **B-D.** Traces of electroretinogram (ERG) recordings from adult eye photoreceptors upon repetitive stimulation with light (left) and quantification of the relative ERG amplitude for indicated genotypes (right). Error bars show mean with SD. N ≥ 6 larvae per genotype. ** P≤0.01, *** P≤0.001. Significance for each genotype was calculated with a T-Test comparing to control flies. **B.** Recordings were taken from 1-3 days post-eclosion animals that were raised in a 12hr light/dark cycle. “On” and “Off” transients indicated by closed and open arrowhead, respectively. **C.** Recordings were taken from 1-3 days post- eclosion animals that were raised in a 24hr dark. **D.** Recordings were taken from four week aged animals that were raised in a 12hr light/dark cycle.

In addition to measuring neuronal activity, we analyzed *dKO* neurons for changes in morphology and molecular markers. Confocal imaging of the NMJ in *dKO* 3^rd^ instar larvae did not reveal obvious changes in synapse morphology or markers of synapse function (Supplemental Figure 4). In contrast, using transmission electron microscopy (TEM) of sectioned adult eyes, we observed reduced photoreceptor number and aberrant morphology such as enlarged photoreceptors and thinner glia in *dKO* animals (Figure 5A-C), suggestive of degeneration. These phenotypes were rescued by a genomic transgene, but not with single ORF rescue constructs (Figure 5A-C, Supplemental Figure 5).

**Figure 5.**
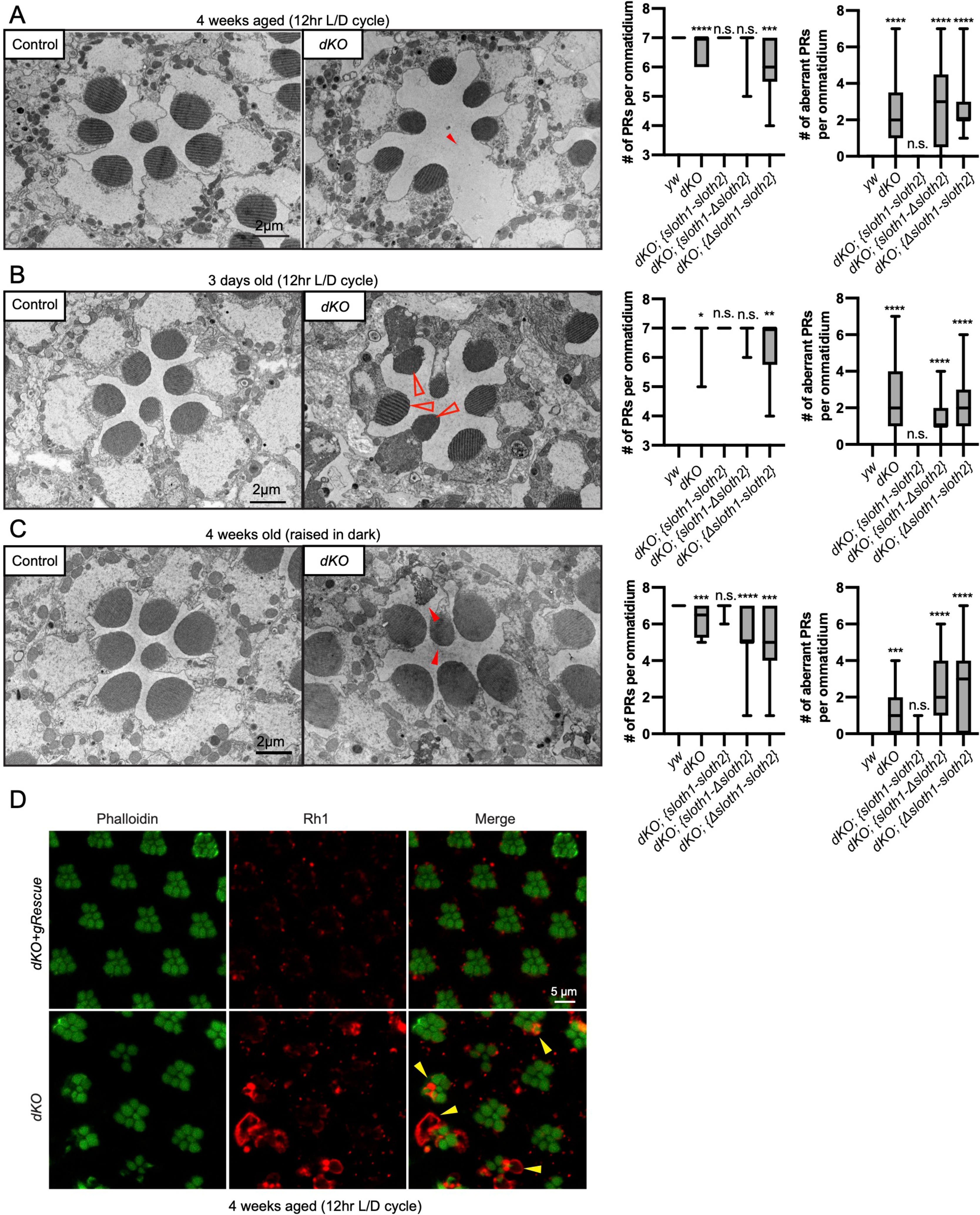
Loss of *sloth1*-*sloth2* causes neurodegeneration. A-C. Transmission electron microscopy (TEM) images of sectioned adult eye photoreceptors (left) and quantification of photoreceptor number and aberrant photoreceptors (right). Scalebar is 2µm. Filled red arrows indicate dead or dying photoreceptors. Open red arrows indicate unhealthy photoreceptors. Error bars show mean with SD. N ≥ 8 ommatidium per genotype. **A.** 4 weeks old raised in a 12hr light/dark cycle. **B.** 3 days old raised in a 12hr light/dark cycle. **C.** 4 weeks old raised in 24hr dark. **D.** Confocal microscopy of adult eye photoreceptors stained with phalloidin (green) and anti-Rh1 (red). Animals were 4 weeks old and raised in a 12hr light/dark cycle. Arrowheads indicate photoreceptors with higher levels of Rh1.

Furthermore, these phenotypes were similar between young and aged flies, as well as aged flies raised in the dark (Figure 5A-C, Supplemental Figure 5). It is known that mutations affecting the turnover of Rhodopsin protein (Rh1) can lead to photoreceptor degeneration (Alloway *et al*. 2000; Jaiswal *et al*. 2015). To test if this mechanism is occurring in *dKO* photoreceptors, we imaged Rh1 protein levels using confocal microscopy. We observed Rh1 accumulation in degenerating *dKO* photoreceptors in 4 week aged flies exposed to light (Figure 5D). However, Rh1 accumulation was milder in 4 week aged flies raised in the dark (Supplemental Figure 6). These results point out that light stimulation, and hence activity, enhance degeneration due to Rh1 accumulation in *dKO* animals.

### Sloth1 and Sloth2 localize to mitochondria and their loss impairs normal respiration and ATP production

Mitochondrial dysfunction in *Drosophila* is known to cause phenotypes that are reminiscent of loss of *sloth1* and *sloth2*, such as pupal lethality, reduced neuronal activity, photoreceptor degeneration, and Rh1 accumulation in photoreceptors (Jaiswal *et al*. 2015). Therefore, we investigated the possible role of Sloth1 and Sloth2 in mitochondria.

Prior to our work, a large-scale study of human protein localization suggested that SMIM4 and C12orf73 localize to mitochondria in cultured cells (Thul *et al*. 2017). SMIM4 has a predicted mitochondrial targeting sequence using MitoFates (Fukasawa *et al*. 2015) (0.842), but C12orf73, Sloth1, and Sloth2 do not (.0016, 0.016, 0.009, respectively). In addition, SMIM4 and Sloth1 are predicted to localize to the mitochondrial inner membrane using DeepMito (0.93 and 0.73, respectively), but C12orf73 and Sloth2 are not (0.66 and 0.49, respectively) (Savojardo *et al*. 2020). To test if Sloth1/2 localize to mitochondria in *Drosophila*, we transfected S2R+ cells with Sloth1-FLAG or Sloth2-FLAG. Both Sloth1 and Sloth2 proteins colocalized with the mitochondrial marker ATP5α (Figure 6A). Furthermore, Sloth1-FLAG and Sloth2-FLAG were enriched in mitochondrial fractions relative to cytoplasmic fractions (Figure 6B). Similar results were observed using stable S2R+ cell lines that express streptavidin binding peptide (SBP) tagged Sloth1 or Sloth2 under a copper inducible promoter (*MT-Sloth1-SBP* and *MT-Sloth2-SBP*) (Figure 6C).

**Figure 6.**
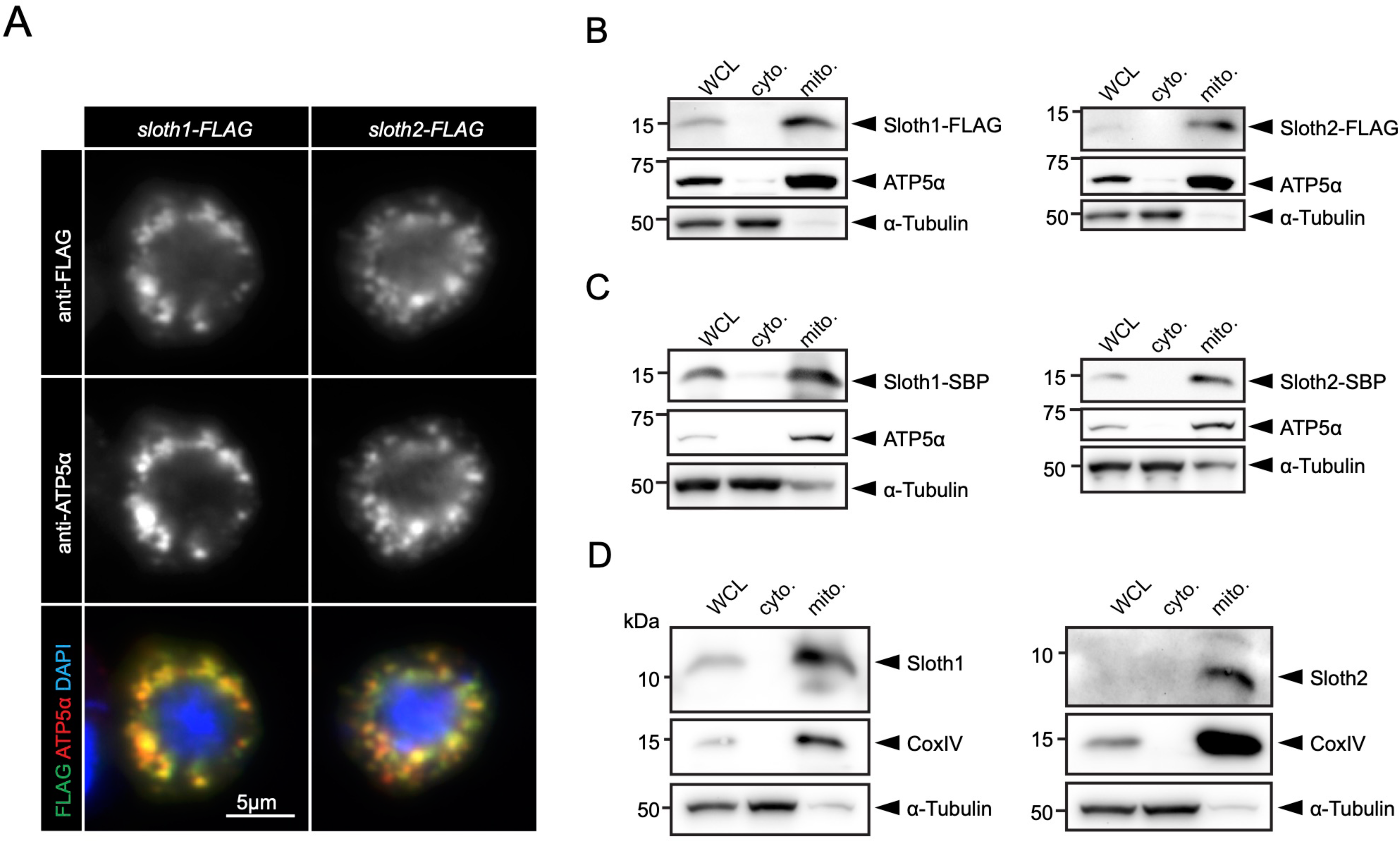
Sloth1 and Sloth2 localize to mitochondria. **A.** Confocal microscopy of S2R+ cells transfected with Sloth1-FLAG or Sloth2-FLAG and stained with anti-FLAG (green) and anti-ATP5alpha (red). DAPI (blue) stains nuclei. **B-D.** SDS-PAGE and western blotting of S2R+ cellular fractions. WCL = Whole Cell Lysate, cyto. = cytoplasmic lysate, mito. = mitochondrial lysate. Mitochondrial control = ATP5alpha, cytoplasmic control = alpha-tubulin. Each lane loaded equal amounts of protein (15µg/lane). Blots were stripped and reprobed after detection of each antigen. **B.** Transfected Sloth1-FLAG or Sloth2-FLAG. **C.** Stable cells expressing copper-inducible Sloth1-SBP or Sloth2-SBP. **D.** Stable cells expressing copper-inducible Sloth1-SBP or Sloth2-SBP.

Next, we raised antibodies to Sloth1/2 to determine their endogenous localization. Using two independently generated antibodies for each peptide, immunolocalization in larval brains from wild-type or *sloth1/2* dKO animals showed no overlapping signal with a mitochondrial marker and no clear signal above background (Supplemental Figure 7). Furthermore, we did not detect Sloth1 or Sloth2 bands of the expected molecular weight on western blots from wild-type S2R+ whole cell lysates or isolated mitochondria using anti-Sloth1, anti- Sloth2, anti-SMIM4, or anti-C12orf73 (Supplemental Figure 8A-C). In contrast, anti-Sloth1 western blots of mitochondria isolated from 3^rd^ instar larvae and adult thoraxes showed a <15kDa band that is absent from *sloth1/2* KO or RNAi samples (Supplemental Figure 8D), suggesting this band corresponds to endogenous Sloth1. Unfortunately, anti-Sloth2 failed to detect a similar band under the same conditions (Supplemental Figure 8D).

Since our Sloth1/2 antibodies may not be sensitive enough to detect the endogenous peptides, we generated a stable S2R+ cell line expressing *sloth1/2* transcript under a copper inducible promoter (*MT-sloth1/2*) and induced expression for 16hrs. Anti-Sloth1 and anti-Sloth2 western blots of mitochondria isolated from *MT-sloth1/2* cells detected <15kDa bands that did not appear in wild-type S2R+ cells, and thus are likely Sloth1 and Sloth2 peptides translated from the overexpressed *sloth1/2* transcript (Supplemental Figure 8B).

Furthermore, Sloth1 and Sloth2 were enriched in *MT-sloth1/2* mitochondrial fractions relative to cytoplasmic fractions (Figure 6D), similar to the results obtained with FLAG and SBP-tagged peptides (Figures 6B-C). Based on their amino acid sequence, Sloth1 and Sloth2 are predicted to run at 9.3kDa and 6.7kDa, respectively. While Sloth1 does appear to run larger than Sloth2, both peptides run ∼2kDa larger than expected (Figure 6D).

A method of assaying defects in mitochondrial function is measuring cellular oxygen consumption from live cells with a Seahorse stress test. Since this typically involves assaying a monolayer of cells, we generated KO S2R+ cell lines using CRISPR/Cas9. Compared to control cells, single KO and double KO S2R+ cells (Supplemental Figure 9A, B) had reduced basal respiration (Figure 7A, B), ATP production (Supplemental Figure 9C), and proton leaks (Supplemental Figure 9D). Results were similar for single KO and dKO lines. These results suggest that both *sloth1* and *sloth2* are required to support normal mitochondrial respiration in S2R+ cells.

**Figure 7.**
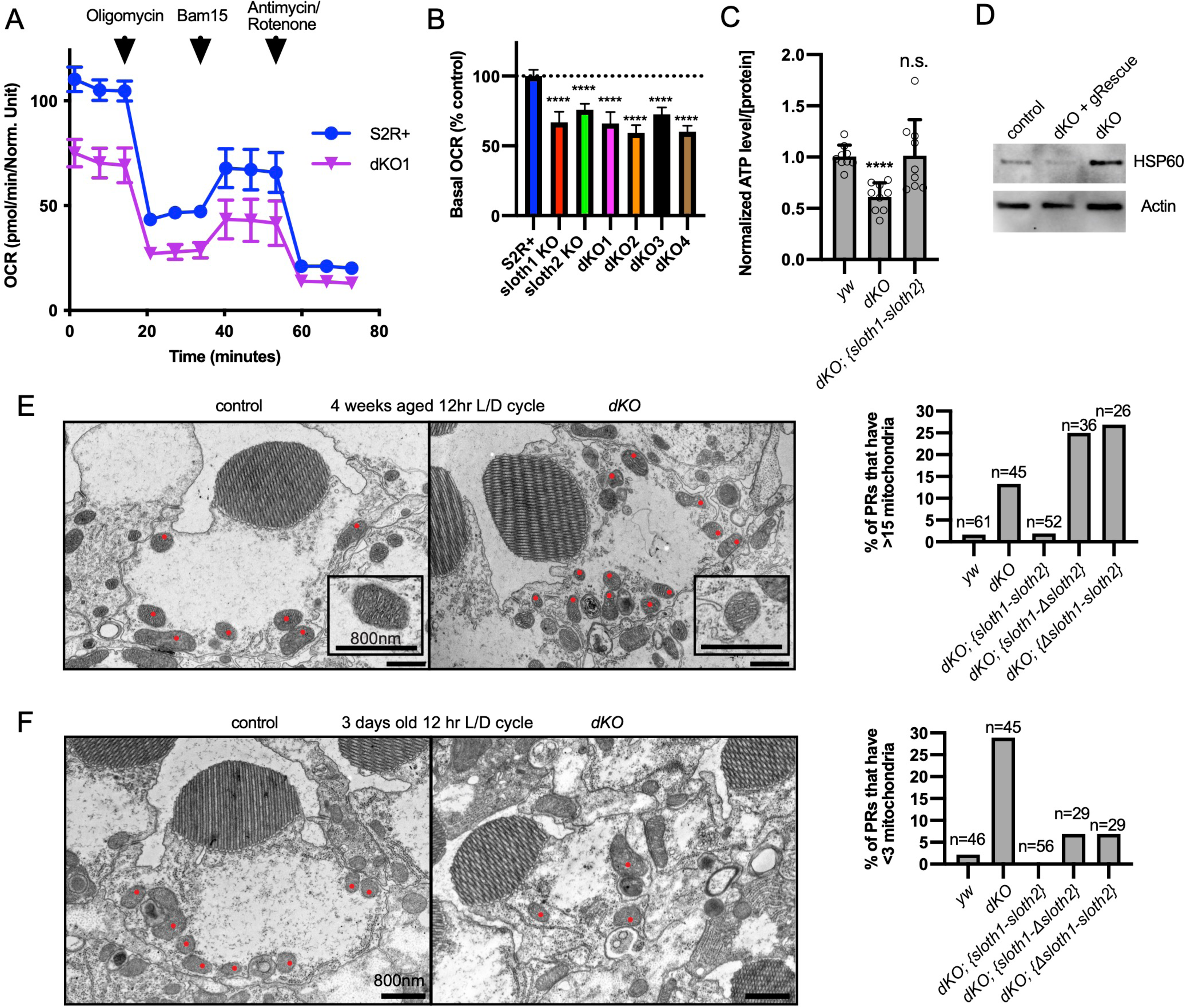
*sloth1*-*sloth2* are important for mitochondrial function. **A.** Seahorse mitochondrial stress report for wildtype S2R+ and dKO #1 cells. Error bars show mean with SD. N=6 for each genotype. **B.** Quantification of basal OCR (timepoint 3) in panel A and including data from single KO and additional dKO cell lines. Significance of KO lines was calculated with a T-test compared to S2R+. Error bars show mean with SD. **** P≤0.0001. N=6 for each genotype. **C.** Quantification of ATP levels in 3^rd^ instar larvae. Error bars show mean with SEM. N = 3 experiments. **D.** Western blot from lysates of 3^rd^ instar larval brains. **E-F.** TEM images of sectioned adult photoreceptors (left) and quantification of mitochondria number (right). Mitochondria are indicated with red dots. Error bars show mean with SD. Sample size indicated on graph. **E.** Adult flies are 4 weeks old and raised in a 12hr light/dark cycle. **F.** Adult flies are 3 days old and raised in a 12hr light/dark cycle.

Next, we assayed *sloth1* and *sloth2* mutant flies for defects in mitochondrial function. ATP levels are an important indicator of mitochondrial function (Kann and Kovacs 2007; Golpich *et al*. 2017) and mutations in *Drosophila* mitochondrial genes can lead to reduced ATP levels (Jaiswal *et al*. 2015).

Indeed, *dKO* larvae had ∼60% ATP compared to control larvae, which was rescued by a genomic transgene (Figure 7C). Impaired mitochondrial function can also lead to cellular stress responses, such as increased expression of the mitochondrial chaperone Hsp60 (Pellegrino *et al*. 2013). Western blot analysis showed that *Drosophila* Hsp60 was elevated in lysates from mutant larval brains compared to control, and this effect was rescued by a genomic transgene (Figure 7D). Finally, mitochondrial dysfunction can cause changes in mitochondrial morphology and number (Trevisan *et al*. 2018). There were no obvious changes in mitochondrial morphology in mutant larval motor neurons (Supplemental Figure 4, Supplemental Figure 9E), and adult mutant photoreceptors contained mitochondria with normal cristae (Figure 7E). In contrast, mitochondrial number was increased in mutant photoreceptors in aged animals (Figure 7E, Supplemental Figure 10A) and decreased in mutant photoreceptors in young animals (Figure 7F, Supplemental Figure 10B). In all, these data suggest that Sloth1 and Sloth2 localize to mitochondria and are important to support respiration and ATP production.

### Sloth1/2 regulate respiratory complex III assembly

While our study was in preparation, two studies demonstrated that human SMIM4 and C12orf73 are inner mitochondrial membrane peptides important for complex III assembly and physically interact with complex III subunits (Zhang *et al*. 2020; Dennerlein *et al*. 2021). If Sloth1 or Sloth2 have similar roles in *Drosophila*, this could explain why *sloth1/2* mutant flies have reduced ATP production.

To test for a role in Sloth1/2 in respiratory complex assembly, we visualized the relative abundance of individual complexes and subunits in wild-type vs *sloth1/2* loss of function animals. First, we resolved native respiratory complexes using blue native polyacrylamide gel electrophoresis (BN-PAGE). Using mitochondria isolated from adult thorax, we identified the five respiratory complexes (CI, CII, CIII, CIV, CV) based on molecular weight and a previous study that established this protocol (Garcia *et al*. 2017). Importantly, a ∼600kDa band corresponding to complex III was diminished in mitochondria isolated from thoraxes with *sloth1/2* knockdown (Figure 8A). Similarly, the complex III band was diminished in mitochondria isolated from *sloth1/2* knockout 3^rd^ instar larvae (Figure 8B). This change was rescued by a wild-type genomic transgene, but not single paralog transgenes (Figure 8B). Next, we detected individual respiratory subunits by SDS-PAGE and western blotting of isolated mitochondria. Using antibodies that recognize UQCR-C2, the fly homolog of human complex III subunit UQCRC2, we found that the ∼40kDa band corresponding to UQCR-C2 was diminished in mitochondria isolated from *sloth1/2* RNAi adult thoraxes (Figure 8C), as well as *sloth1/2* knockout 3^rd^ instar larvae (Figure 8D).

**Figure 8.**
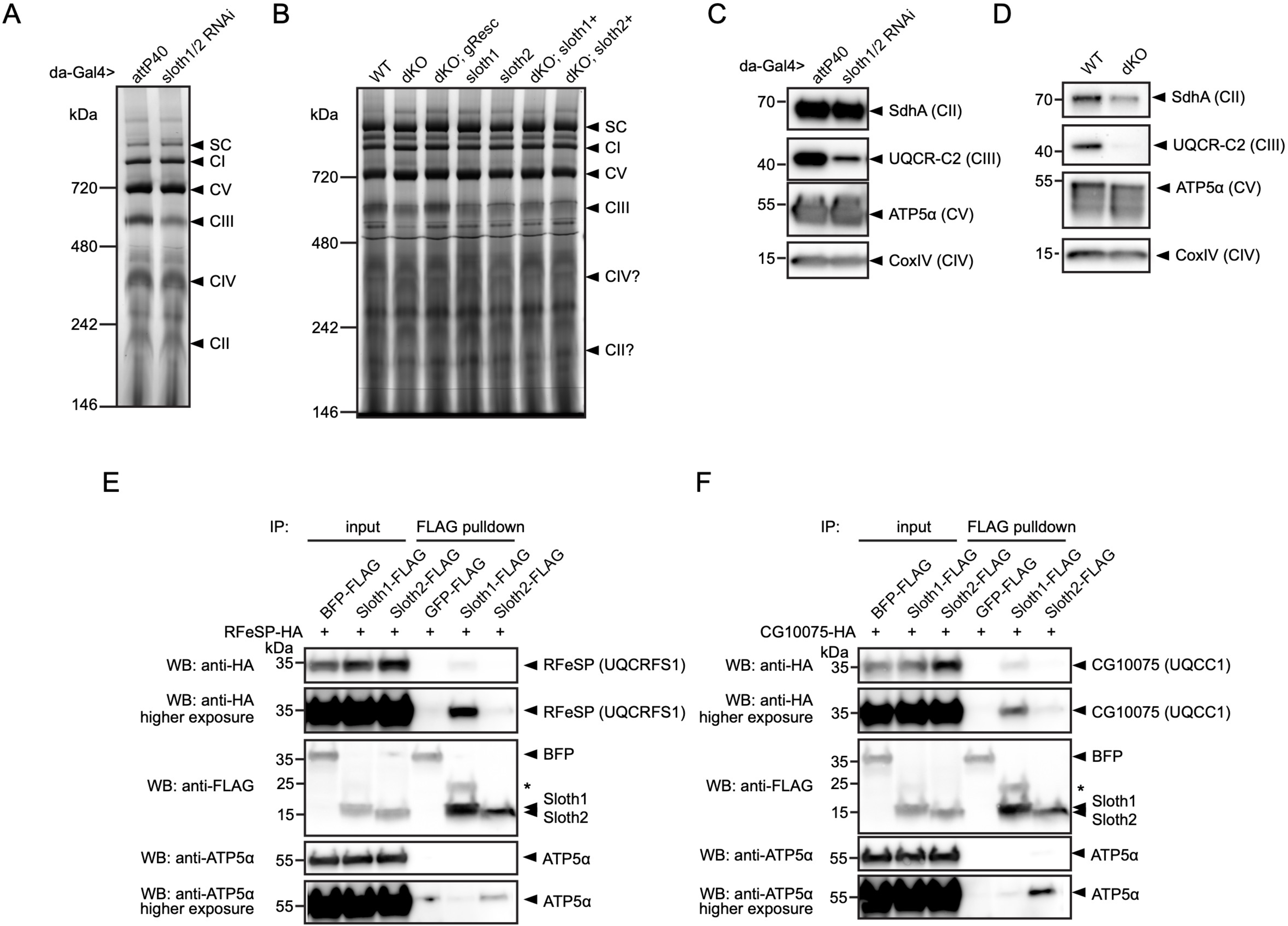
Sloth1 and Sloth2 physically interact with complex III and regulate its assembly. A-B. Blue native PAGE gel of mitochondria isolated from **A.** 10 adult thoraxes and **B.** 10 whole 3^rd^ instar larvae of indicated genotype. Bands corresponding to native respiratory complexes are indicated with arrowheads. **C-D.** SDS-PAGE and western blotting of mitochondria isolated from **C.** adult thorax and **D.** whole 3^rd^ instar larvae of indicated genotype. Each lane loaded equal amount of protein (15µg). Blots were stripped and reprobed after detection of each antigen. **E-F.** Western blots from co-immunoprecipitation experiments in transfected S2R+ cells using Sloth1-FLAG and Sloth2-FLAG as bait and either **E.** RFeSP-HA or **F.** CG10075-HA as prey. Blots were striped and reprobed after detection of each antigen. Arrowheads indicated expected band, asterisks indicate unknown bands.

To test whether Sloth1/2 physically interact with subunits of mitochondrial complex III, we performed co-immunoprecipitation experiments in transfected S2R+ cells. SMIM4 and C12orf73 interact with complex III subunits UQCC1 and UQCRFS1, respectively (Zhang *et al*. 2020; Dennerlein *et al*. 2021). Therefore, we tested if Sloth1 or Sloth2 could immunoprecipitate the fly homologs CG10075 (dUQCC1) or RFeSP (dUQCRFS1). Using Sloth1-FLAG as bait, we detected CG10075-HA (Figure 8E) and RFeSP-HA (Figure 8F) binding to anti-FLAG beads. In contrast, Sloth2-FLAG pulled-down CG10075-HA and RFeSP-HA weakly or was at background levels (Figure 8E,F). Together, these results suggest that Sloth1/2 are required for proper complex III assembly, mediated through physical interaction with complex III subunits.

### Sloth1 and Sloth2 act in a stoichiometric complex

We speculated that Sloth1 and Sloth2 could physically interact, based on the observation that both share the same loss of function phenotypes and subcellular localization. Indeed, some paralogs bind to the same protein complex (Szklarczyk *et al*. 2008) and there is a tendency for proteins in the same complex to be co-expressed (Papp *et al*. 2003). To confirm this putative interaction between Sloth1 and Sloth2, we used co-immunoprecipitation and western blotting. This revealed that Sloth1-FLAG could immunoprecipitate Sloth2-HA (Figure 9A), and reciprocally Sloth2-FLAG (Figure 9B) could immunoprecipitate Sloth1-HA. Interestingly, the levels of tagged peptide in cell lysates were higher when the opposite peptide was overexpressed (Figure 9A,B). Proteins in a complex commonly have important stoichiometry and unbound proteins can be degraded to preserve this balance (Papp *et al*. 2003; Sopko *et al*. 2006; Veitia *et al*. 2008; Prelich 2012; Bergendahl *et al*. 2019). Furthermore, imbalanced protein complex stoichiometry can lead to haploinsufficient or dominant negative phenotypes (Papp *et al*. 2003; Sopko *et al*. 2006; Veitia *et al*. 2008; Prelich 2012; Bergendahl *et al*. 2019).

**Figure 9.**
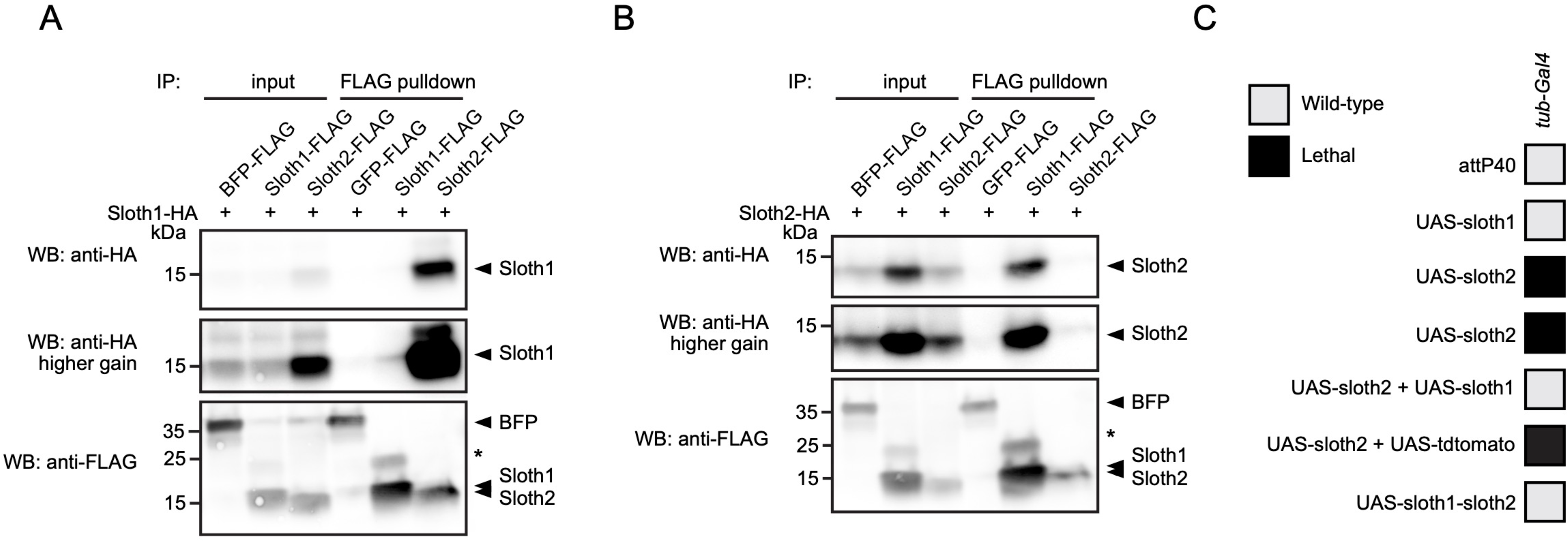
Sloth1 and Sloth2 act in a stoichiometric complex. A-B. Western blots from co-immunoprecipitation experiments in transfected S2R+ cells. **A-B.** Immunoprecipitation using Sloth1-FLAG and Sloth2-FLAG as bait and either **A.** Sloth1-HA or **B.** Sloth2-HA as prey. Blots were striped and reprobed after detection of each antigen. Arrowheads indicated expected band, asterisks indicate unknown bands. **C.** Developmental viability assay using *tub-Gal4* to overexpress indicated transgenes throughout development. Crosses resulting in no viable adults are scored as lethal (black box).

To test this possibility for Sloth1/2, we overexpressed either *sloth1* or *sloth2* in vivo. Low-level ubiquitous overexpression (using *da-Gal4*) of either *UAS-sloth1* or *UAS-sloth2* cDNA had no effect on adult fly viability (Figure 2L). To increase expression levels, we used the strong ubiquitous driver *tub-Gal4*. Whereas *tub>sloth1* flies were viable as adults, *tub>sloth2* animals were 100% pupal lethal (Figure 9C). However, *tub>sloth2* animals could be rescued to adulthood by co-expression of *sloth1*. Importantly, this rescue was not due to dilution of the Gal4 transcription factor on two *UAS* transgenes, since co-expression of *UAS-tdtomato* did not rescue *tub>sloth2* lethality. Finally, *tub-Gal4* overexpression of the entire *sloth1-sloth2* bicistronic transcript resulted in viable adult flies. In all, these results suggest that Sloth1 and Sloth2 interact in a complex where their stoichiometric ratio is important for normal function.

## Discussion

Here, we have assigned new functions to two previously uncharacterized smORF peptides. Sloth1 and Sloth2 appear to be distantly-related paralogs, yet each is important to support mitochondrial and neuronal function in *Drosophila*. We propose a model where Sloth1 and Sloth2 peptides are translated from the same transcript, imported into mitochondria where they interact with each other and complex III to promote its assembly (Figure 10). Our results are supported by two recent studies published during preparation of this manuscript, in which human Sloth1 (SMIM4) and Sloth2 (C12orf73/Brawnin) were discovered as novel mitochondrial complex III assembly factors in cultured human cells and zebrafish (Zhang *et al*. 2020; Dennerlein *et al*. 2021).

**Figure 10.**
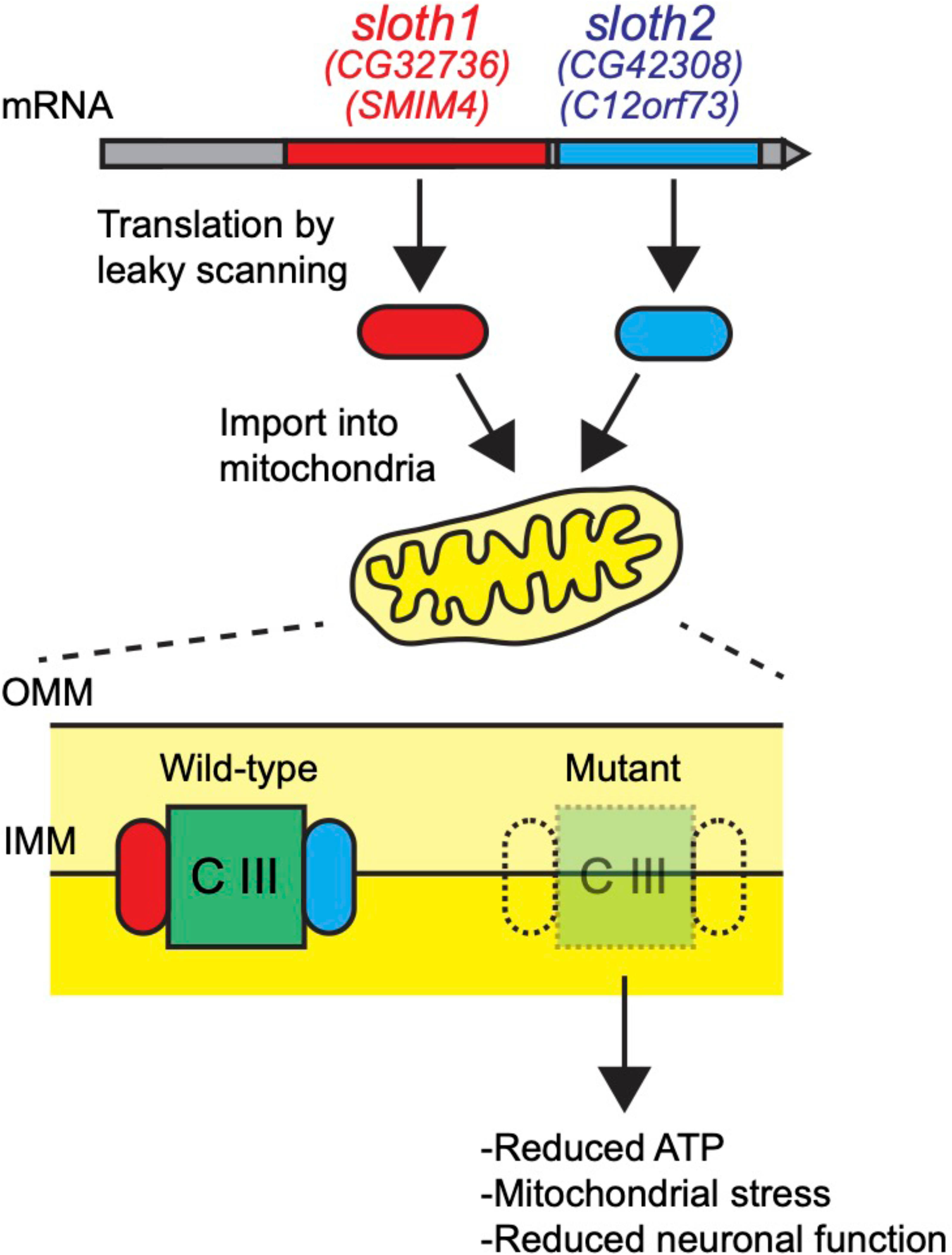
**Model for Sloth1 and Sloth2 bicistronic translation and function in mitochondria**

Muti-cistronic genes are relatively rare in eukaryotes, but some have been characterized in *Drosophila* (Galindo *et al*. 2007; Magny *et al*. 2013) and mammals (Karginov *et al*. 2017). Similar to operons in prokaryotes, it is thought that multicistronic transcripts allow for coordinated expression of proteins in the same pathway or complex (Karginov *et al*. 2017). Indeed, the similarity of loss of function phenotypes between *sloth1* and *sloth2* suggest that they function together in the same pathway/complex. Interestingly, 44/196 annotated bicistronic genes in *Drosophila* contain two ORFs with homology to each other (Flybase, DIOPT), and a recent study suggests that human bicistronic genes containing a smORF frequently encode physically interacting peptide/protein pair (Chen *et al*. 2020). Therefore, related peptides encoded on the same transcript may be a prevalent phenomenon in eukaryotes. ORF translation in multicistronic transcripts can occur by different mechanisms, such as re-initiation of translation, IRES, or leaky ribosome scanning (Van Der Kelen *et al*. 2009). Our data and observations support leaky scanning, and we propose a model whereby both peptides are translated because *sloth1* contains a non-optimal Kozak sequence.

The presence of *sloth1* and *sloth2* orthologs in many eukaryotic species suggest that their function is likely broadly conserved. Indeed, we could rescue the lethality of *sloth1* and *sloth2* mutant flies by expressing their human counterparts. Interestingly, *Plasmodium* and *Arabidopsis* only have homologs with similarity to *sloth2*. Perhaps *sloth2* maintained functions more similar to its common ancestor with *sloth1*. We were unable to identify homologs in some eukaryotes such as yeast, though their amino acid sequence may simply be too diverged for detection using bioinformatic programs such as BLAST.

The physical interactions of Sloth1-Sloth2, Sloth1-RFeSP, and Sloth1-CG10075, and complex III assembly defects in *sloth1/2* loss of function animals, suggest that Sloth1/2 together regulate complex III assembly. Indeed, Sloth1 is bioinformatically predicted to localize to the mitochondrial inner membrane (DeepMito), and Sloth1 and Sloth2 have predicted transmembrane domains (TMHMM 2.0), suggesting they interact with complex III at the inner membrane. This is supported by data showing SMIM4 and C12orf73 are integral membrane proteins in the mitochondrial inner membrane (Zhang *et al*. 2020; Dennerlein *et al*. 2021). In addition, our data suggests that Sloth1 and Sloth2 interact in a stoichiometric manner, explaining why single mutants have the same phenotype as double mutants. This is supported by the finding that SMIM4 protein levels are dependent on the presence of C12orf73 and vice versa (Dennerlein *et al*. 2021).

Perhaps maintenance of the proper ratio of Sloth1/2 is an important factor for optimal complex III assembly. Future experiments could address whether Sloth1 and Sloth2 directly bind each other, or if they require complex III subunits for physical association.

Several observations and experiments suggest that Sloth1/2 peptides do not have equivalent function. The two peptides have weak homology to each other (27% identity) and Sloth1 has 18aa (30%) more than Sloth2, suggesting divergence of function. Unlike Sloth1, Sloth2 does not have a clear mitochondrial-targeting signal. Perhaps Sloth2 has a cryptic signal that is not recognized by prediction software, or Sloth2 may be co-imported with Sloth1. Furthermore, we could not detect robust immunoprecipitation of RFeSP or CG10075 using Sloth2 as bait. Perhaps Sloth2 binds complex III indirectly through Sloth1, or Sloth 2 binds a different complex III subunit. More likely is that both Sloth1 and Sloth2 need to be present for binding to complex III, and the endogenous Sloth1 present under conditions of Sloth2-FLAG overexpression is insufficient for co-IP assays. Sloth2 may also be less stable than Sloth1, which could potentially explain why were unable to detect endogenous Sloth1 using anti-Sloth1 antibodies. Interestingly, only strong overexpression of Sloth2, and not Sloth1, was lethal to flies. Future studies may elucidate the mechanism explaining these functional differences in Sloth1/2.

Neurons have a high metabolic demand and critically depend on ATP generated from mitochondria to support processes such as neurotransmission (Verstreken *et al*. 2005; Kann and Kovacs 2007). Therefore, it is not unexpected that neurodegenerative diseases are frequently associated with mitochondrial dysfunction (Golpich *et al*. 2017). We find similar results in *Drosophila*, where loss of *sloth1* and *sloth2* leads to defects in mitochondrial function, impaired neuronal function, photoreceptor degeneration, and Rh1 accumulation in photoreceptors. Despite finding that the *Gal4-KI* reporter was strongly expressed in neurons and could rescue *sloth1/2* lethality, it is likely these peptides play important roles in other cell types. For example, publicly available RNA-seq data suggest that they are ubiquitously expressed (Flybase). In addition, neuronal expression of *sloth1* or *sloth2* was unable to rescue mutant lethality (Figure 2L). Furthermore, we observed *sloth1/2* loss of function phenotypes in dissected adult thoraxes, which are composed of mostly muscle. At present, there are no reported human disease-associated mutations in *SMIM4* and *C12orf73*. Mutations in these genes might not cause disease, or they might cause lethality. It is also possible that the lack of functional information on these genes has hampered identification of disease-associated mutations.

There is great interest in identifying the complete mitochondrial proteome (Calvo *et al*. 2016), so it is remarkable that Sloth1/2 have been largely missed in proteomic or genetic screens for mitochondrial components. For example, they are not present in bioinformatic and proteomic datasets of fly mitochondrial proteins (Sardiello *et al*. 2003; Chen *et al*. 2015), nor in genetic screens of lethal mutations on the X-chromosome affecting nervous system maintenance (Yamamoto *et al*. 2014). It is possible that the small size of these peptides lead to this discrepancy; due to less frequent mutations in these ORFs, or fewer tryptic products for MS. It is also possible that these peptides form weak interactions with mitochondrial proteins, preventing their immunoprecipitation. Recently, human SMIM4 was identified in a proteomic screen (Dennerlein *et al*. 2021), human C12orf73 was identified in two proteomics screens (Liu *et al*. 2018; Antonicka *et al*. 2020) and a bioinformatic screen (Zhang *et al*. 2020), and mouse SMIM4 was identified in a proteomics screen (Busch *et al*. 2019).

Our discovery of *sloth1* and *sloth2* highlights the effectiveness of loss of function genetics for identifying smORF genes with important biological functions. Recent technical advances such as genome engineering (e.g. CRISPR/Cas9) and massively parallel profiling have the potential to rapidly assign functions to many uncharacterized smORFs (Guo *et al*. 2018; Chen *et al*. 2020). For example, investigation of uncharacterized smORF genes may yield additional important mitochondrial components. Indeed, there is a greater tendency for annotated human smORF peptides to localize to mitochondria (72/719, 10%) compared to the whole proteome (1228/20351, 6%) (UniProt). Interestingly, ∼40 smORF peptides function at the human mitochondrial inner membrane (UniProt), such as the Complex III member UQCRQ (82aa) (Usui *et al*. 1990) and the recently described Mitoregulin/MoxI (56aa) that regulates the electron transport chain and fatty acid β-oxidation (Makarewich *et al*. 2018; Stein *et al*. 2018; Chugunova *et al*. 2019). Therefore, modulation of protein complexes in the inner mitochondrial membrane may be a common function of smORF peptides. As functional annotation of hundreds, perhaps thousands, of smORF genes is becoming easier, many new biological insights are likely to emerge from their analyses.

## Acknowledgements

We thank the TRiP and DRSC for help generating transgenic flies, Dr. Marcia Haigis for use of a Seahorse XF analyzer, Claire Hu, Tera Levin, and Dan Richter for bioinformatics help, Lucy Liu for assistance mounting larvae to image the NMJ, Thai LaGraff for help with qPCR, and Rich Binari and Cathryn Murphy for general assistance. We thank members of the BDGP for discussions. We also thank the HMS MicRoN (Microscopy Resources on the North Quad) Core. J.A.B. was supported by the Damon Runyon Foundation. This work was supported by NIH grants R01GM084947, R01GM067761, R24OD019847, and NHGRI HG009352 (S.E.C). N.P. is an investigator of the Howard Hughes Medical Institute.

## Author Contributions

Conceptualization, J.A.B., B.U., I.P., B.B., S.C., H.B., N.P.; Methodology, J.A.B., B.U., I.P., N.P.; Investigation, J.A.B., B.U., I.P., J.R., F.E., Z.Z.; Writing – Original Draft, J.A.B.; Writing – Review & Editing, J.A.B., B.U., I.P., J.R., F.E., Z.Z., S.C., H.B., N.P.; Supervision, B.B., S.C., D.A.S., H.B., N.P.; Funding acquisition, J.A.B., B.B., S.C., N.P.

## Methods

### Molecular cloning

Plasmid DNAs were constructed and propagated using standard protocols. Briefly, chemically competent TOP10 *E.coli.* (Invitrogen, C404010) were transformed with plasmids containing either Ampicillin or Kanamycin resistance genes and were selected on LB-Agar plates with 100µg/ml Ampicillin or 50µg/ml Kanamycin. Oligo sequences are in Supplemental File 2.

#### *sloth1-sloth2* expression reporters

*pMT-sloth1-RLuc* was constructed by Gibson (NEB E2611) assembly of two DNA fragments with overlapping sequence, 1) 5’UTR, *sloth1* coding sequence, and intervening sequence (*GCAAA*) were amplified from S2R+ genomic DNA. 2) Plasmid backbone was amplified from *pRmHa-3-Renilla* (Zhou *et al*. 2008), which contains a *Metallothionein* promoter and coding sequence for Renilla luciferase. *pMT-sloth1-RLuc* derivatives were constructed by a PCR-based site directed mutagenesis (SDM) strategy.

#### shRNA expression vector for in vivo RNAi

*pValium20-sloth1-sloth2* (aka *UAS-shRNA*, or *JAB200*) was constructed by annealing complementary oligos and ligating into *pValium20* (Ni *et al*. 2011) digested with NheI and EcoRI. See Supplemental Figure 1 for location of target site.

#### sgRNA expression vectors for CRISPR/Cas9

Plasmids encoding two sgRNAs were constructed by PCR amplifying an insert and ligating into *pCFD4* (Port *et al*. 2014) digested with BbsI. sgRNAs constructed: *pCFD4-sloth1* (aka JAB203), *pCFD4-sloth2* (aka GP01169), *pCFD4-sloth1-sloth2* (aka JAB205, for dKO). See Supplemental Figure 1 for location of target sites.

#### Gal4 HDR donor plasmid

*pHD-sloth1-sloth2-Gal4-SV40-loxP-dsRed-loxP* was assembled by digesting *pHD-DsRed-attP* (Gratz *et al*. 2014) with EcoRI/XhoI and Gibson assembling with four PCR amplified fragments: 1) Left homology arm from genomic DNA from *nos-Cas9[attP2]* flies. 2) *Gal4-SV40* from *pAct-FRT-stop-FRT3-FRT-FRT3-Gal4 attB* (Bosch *et al*. 2015). 3) *loxP-dsRed-loxP* from *pHD-DsRed-attP*. 4) Right homology arm from genomic DNA from *nos- Cas9[attP2]* flies.

#### Custom pEntr vectors

Construction of pEntr vectors (for Gateway cloning) was performed by Gibson assembly of PCR amplified backbone from pEntr-dTOPO (Invitrogen C4040-10) and PCR amplified gene coding sequence (when appropriate, with or without stop codon). List of plasmids: *pEntr_sloth1* (from S2R+ cDNA), *pEntr_sloth2* (from S2R+ cDNA), *pEntr_hSMIM4* (from IDT gBlock), *pEntr_hC12orf73* (from IDT gBlock), *pEntr_sloth1-sloth2* transcript (from S2R+ cDNA), *pEntr_sloth1-sloth2* genomic (from S2R+ genomic DNA), and *pEntr_BFP* (from *mTagBFP2*). Derivatives of *pEntr_sloth1-sloth2* genomic that lack *sloth1* or *sloth2* coding sequence, or derivatives of *pEntr_sloth1* or *pEntr_sloth2* with or without only the N-terminal signal sequence, were generated by PCR amplifying the plasmid and reassembling the linearized plasmid (minus the desired sequence) by Gibson.

#### Custom gateway expression vectors

*pMT-GW-SBP* was constructed by digesting *pMK33-SBP-C* (Yang and Veraksa 2017) and *pMK33-GW* (Ram

Viswanatha) with XhoI/SpeI and ligating the GW insert into digested *pMK33- SBP-C* using T4 ligase.

#### Gateway cloning LR reactions

Gateway cloning reactions were performed using LR Clonase II Enzyme mix (Invitrogen 11791-020). See Supplemental File 3 for plasmids constructed by Gateway reactions. Additional plasmids obtained were *pEntr_RFeSP* (DmCD00481962), *pEntr_CG10075* (DmCD00473802) (The FlyBi

Consortium; https://flybi.hms.harvard.edu/), *pAWF* and *pAWH* (Carnegie Science/Murphy lab), *pWalium10-roe* (Perkins *et al*. 2015), and *pBID-G* (Wang *et al*. 2012).

### Fly genetics

Flies were maintained on standard fly food at 25°C. Wild-type (WT) or control flies refers to *yw*. The *yv; attP40* strain is used as a negative control for experiments involving an shRNA or sgRNA transgene inserted into *attP40*.

Fly stocks were obtained from the Perrimon lab collection, Bloomington Stock center (indicated with BL#), or generated in this study (see below). Bloomington Stocks: *yw* (1495), *yv; P{y[+t7.7]=CaryP}attP40* (36304), *yv,P{y[+t7.7]=nos- phiC31\int.NLS}X; P{y[+t7.7]=CaryP}attP40* (25709), *P{y[+t7.7]=nos- phiC31\int.NLS}X, y[1] sc[1] v[1] sev[21]; P{y[+t7.7]=CaryP}attP2* (25710), *w[1118]; Dp(1;3)DC166, PBac{y[+mDint2] w[+mC]=DC166}VK00033* (30299), *y[1] M{w[+mC]=Act5C-Cas9.P}ZH-2A w[*]* (54590), *y[1] sc[*] v[1] sev[21]; P{y[+t7.7] v[+t1.8]=nos-Cas9.R}attP2* (78782), *w[*]; P{w[+mC]=UAS-2xEGFP}AH2* (6874), *w[1118]; P{w[+mC]=UAS-GFP.nls}14* (4775), *y1 w*; P{tubP-GAL4}LL7/TM3, Sb1 Ser1* (5138), *MN-Gal4*, *UAS-mitoGFP* (42737), *MN-Gal4*, *UAS-nSybGFP* (9263), *UAS-tdTomato* (92759), *elav-Gal4* (8760). Perrimon Lab stocks: *w; da-Gal4*, *lethal/FM7-GFP*.

Transgenic flies using PhiC31 integration were made by injecting attB-containing plasmids at 200ng/ul into integrase-expressing embryos that contained an attP landing site (attP40 or attP2). Injected adults were outcrossed to balancer chromosome lines to isolate transgenic founder flies and eventually generate balanced stocks. *pCFD4-sloth1[attP40]* (aka JAB203), *pCFD4-sloth2[attP40]* (aka GP01169), *pCFD4-sloth1-sloth2[attP40]* (aka JAB205, for dKO), *pValium20- sloth1-sloth2[attP40]* (aka *UAS-shRNA*, or *JAB200*) lines were selected with *vermillion+*. *pWalium10-sloth1[attP2]*, *pWalium10-sloth2[attP2]*, *pValium10- sloth2[attP40]*, *pWalium10-hSMIM4[attP2]*, *pWalium10-hC12orf73[attP2]*, *pWalium10-sloth1-sloth2*transcript*[attP2]*, *pBID-{sloth1-sloth2}[attP40]*, *pBID-{Δsloth1-sloth2}[attP40]*, *pBID-{sloth1-Δsloth2}[attP40]* were selected with *white+*.

*sloth1*-KO, *sloth2-KO,* and *dKO* fly lines were made by crossing sgRNA-expressing transgenic lines to *nos-Cas9[attP2]* flies, outcrossing progeny to *FM7- GFP* balancer flies, and screening progeny founder flies for deletions by PCR and Sanger sequencing.

*Gal4-KI* flies were made by injecting sgRNA plasmid (JAB205) and *pHD-sloth1- sloth2-Gal4-SV40-loxP-dsRed-loxP*, each at 200ng/ul, into embryos expressing Cas9 in the germ line (*nos-Cas9*). Injected adults were outcrossed to *FM7-GFP* flies, progeny were screened for RFP+ expression, and RFP+ founder lines were confirmed by PCR for a correct knock-in.

Knockdown crosses were performed by crossing *da-Gal4* with *pValium20-sloth1- sloth2[attP40]*/*CyO* (aka *UAS-shRNA*, or *JAB200*) or *attP40/CyO* as a negative control. Quantification of viability was performed by counting the number of progeny with or without the CyO balancer. A Chi-square test was used to determine if the ratio of non-balancer flies (CyO^-^) to balancer flies (CyO^+^) was significantly altered in shRNA crosses compared to control crosses. Data was analyzed using Excel and Prism.

For climbing assays, *da-Gal4/shRNA or da-Gal4/attP40* adult progeny were aged 1 week after eclosion and 10 flies were transferred into empty plastic vials without use of CO2. Climbing ability was quantified by tapping vials and recording the number of flies that climb to the top of the vial within 10 seconds, using video analysis. Climbing assays with the same 10 flies were performed three times and averaged. Three biological replicates were performed for each genotype. A T-Test was used to calculate statistical significance. Data was analyzed using Excel and Prism.

Somatic knockout crosses were performed by crossing *Act-Cas9* to *sgRNA[attP40]/CyO* or *attP40/CyO* as a negative control. *Act- Cas9/sgRNA[attP40]* female and male progeny were analyzed for phenotypes. Quantification of viability was performed by counting the number of progeny with or without the *CyO* balancer. A Chi-square test was used to determine if the ratio of non-balancer flies (*CyO*^-^) to balancer flies (*CyO*^+^) was significantly altered in somatic knockout crosses compared to control crosses. Male and female progeny were analyzed separately because they differ in the number of copies of the endogenous *sloth1-sloth2* loci on the X-chromosome. Data was analyzed using Excel and Prism.

Mutant and genomic rescue crosses were performed by crossing *mutant/FM7- GFP* females to genomic rescue constructs or *attP40* as a negative control. *mutant/Y* hemizygous male progeny were analyzed for phenotypes.

Quantification of viability was performed by counting the number of *mutant/Y* vs *FM7GFP* male progeny. Gal4/UAS rescue crosses were performed by crossing *mutant/FM7-GFP;; da-Gal4* females to *UAS-X* lines. Additionally, *Gal4-KI/FM7- GFP* females were crossed to *UAS-X*. Rare *sloth1*-KO, *sloth2-KO, dKO,* and *Gal4-KI* hemizygous adult males normally die by sticking to the fly food after they eclose. To collect these rare mutants for further analysis (scutellar bristle images, climbing assays), we inverted progeny vials so that mutant adults fell onto the dry cotton plug once they eclose.

Overexpression crosses were performed by crossing *tub-Gal4/TM3* females to *UAS-X* lines. At least 100 *tub-Gal4/UAS-X* progeny were analyzed for phenotypes.

### Cell fractionation and mitochondrial isolation

To isolate mitochondria from S2R+ cells, cell pellets were resuspended in 1.1ml hypotonic buffer (10 mM NaCl, 1.5 mM MgCl2, 10 mM Tris- HCl pH 7.5), transferred to cold glass dounce on ice, and incubated for 10min to induce cell swelling. Cells were homogenized with 10 strokes using pestle B (tight pestle), followed by addition of 800µl of 2.5x homogenization buffer (525mM mannitol, 175 mM sucrose, 12.5 mM Tris-HCl pH 7.5 and 2.5 mM EDTA). Homogenates at this step are considered whole cell lysate (WCL). WCL was centrifuged at 1,300g for 5min at 4°C, supernatant transferred to a new tube, repeated centrifugation.

Supernatant was transferred to a new tube and centrifuged at 17,000g for 15min at 4°C. Supernatant was removed (cytoplasmic fraction) and 2ml 1x Homogenization buffer (210 mM mannitol, 70 mM sucrose, 5 mM Tris-HCl pH 7.5 and 1 mM EDTA) was added to the pellet. The centrifugation was repeated and 250µl 1x Homogenization buffer was added to the pellet (mitochondrial fraction). For SDS-PAGE comparisons of cell fractions, WCL, cytoplasmic, and mitochondria were lysed in RIPA buffer and protein concentration normalized by BCA assay (Thermo Fischer, 23227).

Mitochondrial isolation from 7 day old adult thoraxes and whole 3^rd^ instar larvae was modified from (Garcia *et al*. 2017). Briefly, dissected adult male thoraxes or whole 3^rd^ instar male larvae were placed into 100µl mitochondrial isolation buffer (250mM Sucrose, 150mM MgCl2, 10mM Tris-HCl pH 7.4) on ice. Thoraxes were ground using a blue pestle and a motorized pestle holder. 400µl mitochondrial isolation buffer was added to homogenized thoraxes and samples were centrifuged at 500g at 4°C for 5min to pellet debris and tissues. Supernatant was transferred to a new tube and the centrifugation repeated. Supernatant was transferred to a new tube and centrifuged at 5000g at 4°C for 5min to pellet mitochondria. The mitochondrial pellet was washed 2x by adding 1ml mitochondrial isolation buffer and repeating centrifugation at 5000g at 4°C for 5min. For BN-PAGE experiments, 10 thoraxes or 10 whole 3rd instar larvae were used. For SDS-PAGE, 30 thoraxes or 30 whole 3^rd^ instar larvae were used, and mitochondria were lysed in RIPA buffer and protein concentration normalized by BCA assay (Thermo Fischer, 23227).

### Blue Native PAGE (BN-PAGE) of mitochondrial respiratory complexes

Native mitochondrial respiratory complexes were visualized by Blue Native PAGE (BN-PAGE) gels following the manufacturer’s instructions protocols (Nativepage 12% Bis Tris Protein Gels, 1.0 mm, 15 well, Thermo Fisher Scientific BN1003BOX). Mitochondrial pellets from 10 thoraxes or 10 larvae were resuspended in 20ul sample buffer cocktail (5µl sample buffer, 8µl 5% digitonin, 7µl H20, 2µl 5% Coomassie G-250 sample additive). 15µl sample ran on each lane.

### Cell culture

*Drosophila* S2R+ cells (Yanagawa *et al*. 1998), or S2R+ cells stably expressing Cas9 and a mCherry protein trap in *Clic* (known as PT5/Cas9) (Viswanatha *et al*. 2018), were cultured at 25°C using Schneider’s media (21720-024, ThermoFisher) with 10% FBS (A3912, Sigma) and 50 U/ml penicillin strep (15070-063, ThermoFisher). S2R+ cells were transfected using Effectene (301427, Qiagen) following the manufacturer’s instructions.

For generating stable cell lines MT-Sloth1-SBP, MT-Sloth2-SBP, and MT- Sloth1/2, S2R+ cells were seeded in 6-well plates and transfected with *pMK33* expression plasmids (see Supplemental File 3). *pMK33* derived plasmids contain a Hygromycin resistance gene and a *Metallothionein* promoter to induce gene expression. After 4 days, transfected cells were selected with 200µg/ml Hygromycin in Schneider’s medium for approximately 1 month. For induction of gene expression, cells were cultured with 500 µM CuSO4 in Schneider’s medium for 16hrs.

For generating KO cell lines, S2R+Cas9 cells were transfected with *tub-GFP* plasmid (gift of Steve Cohen) and an sgRNA-expressing plasmid (*pCFD4-*

*sloth1[attP40]* (aka JAB203), *pCFD4-sloth2[attP40]* (aka GP01169), or *pCFD4- sloth1-sloth2[attP40]* (aka JAB205, for dKO)). 48hrs after transfection, cells were resuspended in fresh media, triturated to break up cell clumps, and pipetted into a cell straining FACS tube (352235 Corning). Single GFP+ cells were sorted into single wells of a 96 well plate containing 50% conditioned media using an Aria- 594 instrument at the Harvard Medical School Division of Immunology’s Flow Cytometry Facility. Once colonies were visible by eye (3-4 weeks), they were expanded and analyzed by PCR and Sanger sequencing.

For co-immunoprecipitation experiments, S2R+ cells were transfected in 100mm petri dishes. Four days after transfection, cells were resuspended and centrifuged at 1000g for 10min at 4°C. Cell pellets were washed once with ice-cold 1x PBS, re-centrifuged, and flash frozen in liquid nitrogen. Cell pellets were subjected to mitochondrial isolation (described above) and mitochondrial pellets were flash frozen in liquid nitrogen. Mitochondrial pellets were resuspended in 250µl mitochondrial lysis buffer (∼.5-1ug/ul final protein concentration), incubated on ice for 30min and centrifuged at 13,000g for 10min at 4°C. The supernatant was incubated with 20µl magnetic anti-FLAG beads (Sigma-Aldrich M8823) for 2hr at 4°C with gentle rocking. Beads were washed 3x in mitochondrial lysis buffer using a magnetic stand and eluted for 30min at 4°C with 20ul 3xFLAG peptide diluted at 1mg/ml in mitochondrial lysis buffer. Mitochondrial lysis buffer: 50 mM Tris-HCl pH 7.4, 150 mM NaCl, 10% glycerol (v/v), 20 mM MgCl2, 1% digitonin (v/w) (Sigma D141), protease inhibitor (Pierce 87786), and 2 mM PMSF added immediately before use.

To measure mitochondrial respiration in S2R+ cells, we performed a Mito Stress Test on a Seahorse XFe96 Analyzer (Agilent, 103015-100). 50,000 cells were seeded into Seahorse XF96 tissue culture microplates and incubated at 25°C overnight. 1hr before analysis, cell culture media was replaced with serum-free Schneider’s media and drugs were loaded into the Seahorse XFe96 Sensor Cartridge (Final concentrations: Oligomycin 1µM, Bam15 .5µM, 1µM Antimyzin/Rotenone “R/A”). Seahorse analysis was performed at room temperature. Mitochondrial respiration recordings were normalized to cell number using CyQUANT (Thermo Fisher C7026) fluorescence on a plate reader. Data analysis was performed using Seahorse Wave Desktop Software 2.6, Excel, and Prism. N=6 wells for each condition. Significance was calculated using a T-Test.

To measure *MT-sloth1-RLuc* reporter expression, S2R+ cells were transfected in white opaque-bottom 96 well plates with *MT-sloth1-RLuc* (or derivatives) and *MT-FLuc* (Firefly Luciferase) (Zhou *et al*. 2008) as an internal control. Briefly, to each well, 10ng of plasmid mix was added, then 10µl Enhancer mix (.8µl Enhancer + 9.2µl EC buffer), and was incubated for 2-5min at room temperature. 20µl of Effectene mix (2.5µl Effectene + 17.5µl EC buffer) was added and incubated for 5-10min at room temperature. 150µl of S2R+ cells (at 3.3x10^5 cells/ml) was added gently to each well and incubated at 25°C. After 3 days incubation, 200µM CuSO4 was added. After 24 hours incubation, media was gently removed from the wells by pipetting and cell luminescence was measured using the Dual-Glo assay (Promega E2920). Two luminescence normalizations were performed. First, for each sample, Renilla luminescence was normalized to Firefly luminescence (Rluc/Fluc). Next, Rluc/Fluc ratios for each sample were normalized to Rluc/Fluc ratios for wild-type *MT-sloth1-RLuc* (aka fold change Rluc/Fluc to WT). For each genotype, N=4. Significance was calculated using a T-test. Data was analyzed using Excel and Prism.

### Western blotting

Protein or cell samples were denatured in 2x SDS Sample buffer (100mM Tris-CL pH 6.8, 4% SDS, .2% bromophenol blue, 20% glycerol, .58 M β- mercaptoethanol) by boiling for 10 min. For western blots using glycine-based gels (Figure 7D, Figure 8C-F, Figure 9A-B, Supplemental Figure 8A,B,D), denatured proteins and Pageruler Prestained Protein Ladder (Thermo Fisher Scientific 26616) were loaded into 4–20% Mini-PROTEAN TGX gels (Biorad 4561096) using running buffer (25 mM Tris, 192 mM glycine, 0.1% SDS, pH 8.3). For western blots using tricine-based gels (Figure 6B-D, Supplemental Figure 8C) (to improve resolution of small peptides), denatured proteins and Precision Plus Protein™ Dual Xtra Prestained Protein Standards (Biorad 1610377) were loaded into 16.5% Mini-PROTEAN® Tris-Tricine Gels (Biorad 4563066) using Tris/Tricine/SDS Running buffer (Biorad 1610744). Gels were ran at 100-200V in a Mini-PROTEAN Tetra Vertical Electrophoresis Cell (Biorad 1658004). Proteins were transferred to Immobilon-FL PVDF (Millipore IPFL00010) in transfer buffer (25 mM Tris, 192 mM glycine) using a Trans-Blot Turbo Transfer System (Biorad 1704150) (Standard SD program). Resulting blots were incubated in TBST (1x TBS + .1% Tween20) for 20min on an orbital shaker, blocked in 5% non-fat milk in TBST for 1 hour at room temperature, and incubated with primary antibody diluted in blocking solution overnight at 4°C. Blots were washed with TBST and incubated in secondary antibody in blocking solution for 4 hours at room temperature. Blots were washed in TBST before detection of proteins. HRP-conjugated secondary antibodies were visualized using ECL (34580, ThermoFisher). Blots were imaged on a ChemiDoc MP Imaging System (BioRad). Antibody complexes were reprobed by incubating blots with stripping buffer (Thermo Scientific 46430) following the manufacturer’s instructions, re-blocked in 5% non-fat milk in TBST, and incubated with primary antibody overnight as described.

For western blots from larval brains, 3^rd^ instar larval brains were dissected in ice cold PBS buffer with protease and phosphatase inhibitors. 10 brains per genotype were homogenized in RIPA buffer and protein concentration was measured by BCA assay (Thermo Fischer, 23227). Equal amounts of protein samples were mixed with 1X Sample buffer (BioRad, 161-0747), boiled for 5 min, and loaded into 4-20% Mini-PROTEAN® TGX gel (Bio-Rad). Gels were then transferred to nitrocellulose membranes using Bio-Rad Trans-Blot SD Semi-Dry Transfer system. Western blots using anti-Hsp60 likely recognize Hsp60A, as opposed to Hsp60B/C/D, because only Hsp60A is expressed in the larval brain (flyrnai.org/tools/dget/web).

Commercially available or published antibodies used for western blotting: rat anti-HA (1:2000, Roche 11867423001) (Figure 9A,B), chicken anti-HA (1:1000, ET- HA100, Aves) (Figure 8E,F), mouse anti-FLAG (1:1000, Sigma F1804), mouse anti-SBP (1:1000, Santa Cruz sc-101595), mouse anti-a-Tubulin (1:20000, Sigma T5168), rabbit anti-GFP (1:5000, Invitrogen A-6455), rabbit anti-Hsp60 antibody (Abcam ab46798), mouse anti-actin (MP Biomedicals 08691002), anti-actin Rhodamine (Biorad 12004163), rabbit anti-SMIM4 (1:10,000, HPA047771), anti-UQCR-C2 (1:1000, (Murari *et al*. 2020)), anti-SdhA (1:1000, (Murari *et al*. 2020)), rabbit anti-C12orf73 (1:1000, HPA038883), anti-mouse HRP (1:3000, NXA931, Amersham), anti-rat HRP (1:3000, Jackson 112-035-062), anti-rabbit HRP (1:3000, Amersham NA934), anti-chicken HRP (1:1000, Sigma SAB3700199), anti-mouse 800 (only used in Figure 8E,F to detect mouse anti-FLAG) (1:5000, Invitrogen A32730). Anti-Sloth1 and Anti-Sloth2 antibodies (1:1000) were raised in rabbits (Genscript, PolyExpress Silver Package).

Epitopes used: Anti-Sloth1 #1: RRLLDSWPGKKRFGC, Anti-Sloth1 #2: CEQQHLQARAANNTN, Anti-Sloth2 #1: CHSTQVDPTAKPPES, Anti-Sloth2 #2: CYKPLEDLRVYIEQE

### Molecular biology

S2R+ cell genomic DNA was isolated using QuickExtract (QE09050, Lucigen). Fly genomic DNA was isolated by grinding a single fly in 50µl squishing buffer (10 mM Tris-Cl pH 8.2, 1 mM EDTA, 25 mM NaCl) with 200µg/ml Proteinase K (3115879001, Roche), incubating at 37°C for 30 min, and 95°C for 2 minutes. PCR was performed using Taq polymerase (TAKR001C, ClonTech) when running DNA fragments on a gel, and Phusion polymerase (M-0530, NEB) was used when DNA fragments were sequenced or used for molecular cloning. DNA fragments were run on a 1% agarose gel for imaging or purified on QIAquick columns (28115, Qiagen) for sequencing analysis. Sanger sequencing was performed at the DF/HCC DNA Resource Core facility and chromatograms were analyzed using Lasergene 13 software (DNASTAR).

For RT-qPCR analysis of *sloth1-sloth2* RNAi knockdown, *da-Gal4* was crossed with *attP40* or *UAS-shRNA* and ten 3^rd^ instar larvae progeny of each genotype were flash frozen in liquid nitrogen. Frozen larvae were homogenized in 600µl Trizol (Invitrogen 15596026) and RNA extracted using a Direct-zol RNA Miniprep kit (Zymo Research, R2050). cDNA was generated using the iScript Reverse Transcription Supermix (BioRad 1708840). cDNA was analyzed by RT-qPCR using iQ SYBR Green Supermix (BioRad 170-8880). qPCR primer sequences are listed in Supplemental File 2. Each qPCR reaction was performed with two biological replicates, with three technical replicates each. Data was analyzed using Bio-Rad CFX Manager, Excel, and Prism. Data from *sloth1-sloth2* specific primers were normalized to primers that amplify *GAPDH* and *Rp49*. Statistical significance was calculated using a T-Test.

### Bioinformatic analysis

Protein similarity between fly and human Sloth1 and Sloth2 orthologs was determined using BLASTP (blast.ncbi.nlm.nih.gov) by defining the percent amino acid identity between all four comparisons. Homologs in other organisms and their gene structure were identified using a combination of BLASTP, Ensembl (www.ensembl.org), HomoloGene (www.ncbi.nlm.nih.gov/homologene), and DIOPT (www.flyrnai.org/diopt). Protein accession numbers: Human *SMIM4* NP_001118239.1, Human *C12orf73* NP_001129042.1, Mouse *SMIM4* NP_001295020.1, Mouse *C12orf73 homolog* NP_001129039.1, Zebrafish *SMIM4* NP_001289975.1, Zebrafish *C12orf73 homolog* NP_001129045.1, Lamprey *SMIM4* XP_032827557.1, Lamprey *C12orf73 homolog* XP_032827559.1, *D.melanogaster CG32736* NP_727152.1, *D.melanogaster CG42308* NP_001138171.1, *Arabidopsis AT5G57080* NP_200518.1, *Arabidopsis AT4G26055* NP_001119059.1, *Plasmodium PF3D7_0709800* XP_002808771.1, Choanoflagellate (*Salpingoeca urceolata*) *m.92763* (Richter *et al*. 2018), Choanoflagellate (*Salpingoeca urceolata*) *sloth2* homolog is unannotated but present in comp15074_c0_seq2 (Richter *et al*. 2018). Sea squirt (*C. intestinalis*) *sloth1* and *sloth2* homologs are unannotated but present in *LOC100183920* XM_018812254.2. Genomic sequences for *sloth1/2* ORFs in *D.melanogaster*, Lamprey, Choanoflagellate, and Sea squirt are shown in Supplemental File 1.

Amino acid sequence of fly and human Sloth1*/*Sloth2 were analyzed for predicted domains using the following programs: MitoFates (http://mitf.cbrc.jp/MitoFates/cgi-bin/top.cgi), DeepMito (http://busca.biocomp.unibo.it/deepmito/), TMHMM 2.0 (http://www.cbs.dtu.dk/services/TMHMM/).

Amino acid sequences were aligned using Clustal Omega (https://www.ebi.ac.uk/Tools/msa/clustalo/) and visualized using Jalview (https://www.jalview.org/).

### Imaging

For imaging adult scutellar bristles, adult flies were frozen overnight and dissected to remove their legs and abdomen. Dissected adults were arranged on a white surface and a focal stack was taken using a Zeiss Axio Zoom V16. Focal stacks were merged using Helicon Focus 6.2.2.

For imaging larval brains, wandering 3^rd^ instar larvae were dissected in PBS and carcasses were fixed in 4% paraformaldehyde for 20min. Fixed carcasses were either mounted on slides in mounting medium (see below), or permeabilized in PBT, blocked for 1hr in 5% normal goat serum (S-1000, Vector Labs) at room temperature, and incubated with primary antibody (anti-Elav) overnight at 4°C, washed with PBT, incubated with secondary antibody (anti-mouse 633) for 4hr at room temperature, washed with PBT and PBS, and incubated in mounting media (90% glycerol + 10% PBS) overnight at 4°C. Larval brains were dissected from carcasses and mounted on a glass slide under a coverslip using vectashield (H- 1000, Vector Laboratories Inc.). Images of larval brains were acquired on a Zeiss Axio Zoom V16 or a Zeiss 780 confocal microscope. Images were processed using Fiji software.

For imaging the larval NMJ, wandering 3^rd^ instar larvae were dissected as previously described (Brent *et al*. 2009). Briefly, larvae were pinned to a Sylgard-coated (Dow 4019862) petri dish, an incision was made along their dorsal surface, their cuticle was pinned down to flatten the body wall muscles, and were fixed in 4% paraformaldehyde for 20min. Fixed carcasses were permeabilized in PBT, blocked for 1hr in 5% normal goat serum (S-1000, Vector Labs) at room temperature, and incubated with primary antibody overnight at 4°C, washed with PBT, incubated with secondary antibody for 4hr at room temperature, washed with PBT and PBS, and incubated in mounting media (90% glycerol + 10% PBS) overnight at 4°C. Whole carcasses mounted on a glass slide under a coverslip using vectashield (H-1000, Vector Laboratories Inc.).

Images of the NMJ were acquired on a Zeiss Axio Zoom V16 or a Zeiss 780 confocal microscope. Images were taken from muscle 6/7 segment A2. Images were processed using Fiji software. Quantification of bouton number from NMJ stained with anti-HRP and anti-Dlg1 was performed by manual counting of boutons in an entire NMJ for wild-type (N=8) and dKO animals (N=7). A T-test was used to determine significance.

For imaging whole larvae, wandering 3^rd^ instar larvae were washed with PBS and heat-killed for 5min on a hot slide warmer to stop movement. Larvae were imaged using a Zeiss Axio Zoom V16 fluorescence microscope.

For imaging the adult brain, ∼1 week old adult flies were dissected in PBS and whole brains were fixed in 4% paraformaldehyde for 20min. Fixed brains were permeabilized in PBT, blocked for 1hr in 5% normal goat serum (S-1000, Vector Labs) at room temperature, incubated with anti-HRP 647 overnight at 4°C, washed with PBT and PBS, and incubated in mounting media (90% glycerol + 10% PBS) overnight at 4°C. Adult brains were mounted on glass slides under a coverslip using vectashield (H-1000, Vector Laboratories Inc.). Images of adult brains were acquired on a Zeiss 780 confocal microscope. Images were processed using Fiji software.

For confocal microscopy of adult photoreceptors, the proboscis was removed and the head was pre-fixed with 4% formaldehyde in PBS for 30 min. After pre-fixation, eyes were removed from the head and fixed an additional 15 minutes. Fixed eyes were washed with PBS 3x for 10 min each and permeabilized in 0.3% Triton X-100 in PBS for 15 min. Permeabilized, fixed samples were blocked in 1X PBS containing 5% normal goat serum (NGS) and 0.1% Triton X-100 for 1 h (PBT). Samples were incubated in primary antibody diluted in PBT overnight at 4°C, washed 3x with PBT, and incubated in secondary antibodies in NGS for 1hr at room temp the next day. Following secondary antibody incubation, samples were washed with PBS and were mounted on microscope slides using vectashield. Samples were imaged with LSM710 confocal with 63X objective and processed using Fiji software.

S2R+ cells transfected with Sloth1-FLAG or Sloth2-FLAG were plated into wells of a glass-bottom 384 well plate (6007558, PerkinElmer) and allowed to adhere for 2 hours. Cells were fixed by incubating with 4% paraformaldehyde for 30min, washed with PBS with .1% TritonX-100 (PBT) 3x 5min each, blocked in 5% Normal Goat Serum (NGS) in PBT for 1hr at room temperature, and incubated in primary antibodies diluted in PBT-NGS overnight at 4°C on a rocker. Wells were washed in PBT, incubated with secondary antibodies and DAPI and washed in PBS. Plates were imaged on an IN Cell Analyzer 6000 (GE) using a 20x or 60x objective. Images were processed using Fiji software.

List of antibodies and chemicals used for tissue staining: rat anti-Elav (1:50, DSHB, 7E8A10), goat anti-HRP 647 (1:400, Jackson Immunoresearch, 123-605-021), mouse anti-ATP5α (1:500, Abcam, ab14748), DAPI (1:1000, Thermo Fisher, D1306), rabbit anti-FLAG (1:1000, Sigma, F7425), mouse anti-FasII (1:25, DSHB, 1D4), mouse anti-brp (1:25, DSHB, nc82), mouse anti-Dlg1 (1:250, DSHB, 4F3), anti-mouse 633 (1:500, A-21052, Molecular Probes), mouse monoclonal anti-Rh1 (1:50, DSHB 4C5), Phalloidin conjugated with

Alexa 488 (1:250, Invitrogen A12379).

### Transmission electron microscopy (TEM) of adult photoreceptors

TEM of *Drosophila* adult retinae were performed following standard electron microscopy procedures using a Ted Pella Bio Wave processing microwave with vacuum attachments. Briefly, whole heads were dissected in accordance to preserve the brain tissue. The tissue was covered in 2% paraformaldehyde, 2.5% Glutaraldehyde, in 0.1 M Sodium Cacodylate buffer at pH 7.2. After dissection, the heads were incubated for 48hrs in the fixative on a rotator at 4°C. The pre- fixed heads were washed with 3X millipore water followed by secondary fixation with 1% aqueous osmium tetroxide, and rinsed again 3X with millipore water. To dehydrate the samples, concentrations from 25%–100% of Ethanol were used, followed by Propylene Oxide (PO) incubation. Dehydrated samples are infiltrated with gradual resin:PO concentrations followed by overnight infiltration with pure resin. The samples were embedded into flat silicone molds and cured in the oven at 62°C for 3-5 days, depending on the atmospheric humidity. The polymerized samples were thin-sectioned at 48-50 nm and stained with 1% uranyl acetate for 14 minutes followed by 2.5% lead citrate for two minutes before TEM examination. Retina were viewed in a JEOL JEM 1010 transmission electron microscope at 80kV. Images were captured using an AMT XR-16 mid-mount 16 mega-pixel digital camera in Sigma mode. Three animals per genotype per condition were used for TEM. At least 30 photoreceptors were used for organelle quantifications. Quantification of photoreceptor number, number of aberrant photoreceptors, and number of mitochondria per photoreceptor, was performed in Prism. Significance was calculated using a T-Test.

### Electrical recordings

#### Intracellular Recording from Larval NMJ

3^rd^ instar larval NMJ recordings were performed as described previously (Ugur et al. 2017). Briefly, free moving larvae are dissected in HL3.1 buffer without Ca^2+^. Recordings were performed by stimulating the segmental nerve innervating a hemisegment A3, Muscle 6/7 through a glass capillary electrode filled with HL3.1 with 0.75 mM Ca^2+^. There were no differences in input resistance, time constant τ, and resting membrane potential among different genotypes tested.

Repetitive stimulations were performed at 10Hz and were reported relative to the first excitatory junction potential (EJP). Data were processed with Mini Analysis Program by Synaptosoft, Clampfit, and Excel. At least 5 animals were used per each genotype per essay. Significance was calculated using a T-Test.

#### Electroretinograms (ERGs)

ERGs were recorded according to (Jaiswal et al. 2015). Briefly, flies were immobilized on a glass slide with glue. Glass recording electrodes, filled with 100 mM NaCl, were placed on the surface of the eye to record field potential. Another electrode placed on the humerals served as a grounding electrode. Before recording ERGs, flies were adjusted to darkness for three minutes. Their response to light was measured in 1sec. intervals for 30 sec. To test if the flies can recover from repetitive stimulation, we recorded ERGs after 30 sec. and 1min constant darkness following repetitive stimulation. Data were processed with AXON-pCLAMP8.1. At least 6 animals were used per each genotype per essay. Significance was calculated using a T-Test.

#### Measurement of ATP levels from larvae

Ten 3^rd^ instar larvae were snap frozen with liquid nitrogen in a 1.5 mL centrifuge tube. Following freezing, samples were homogenized in 100 µl of 6 M guanidine-HCl in extraction buffer (100 mM Tris and 4 mM EDTA, pH 7.8) to inhibit ATPases, and boiled for 3 min. The samples were centrifuged to remove cuticle. Supernatant was serially diluted with extraction buffer and protein concentration was measured using a BCA kit (Thermo Fischer, 23227). For each genotype, ATP levels were measured from equal protein amounts using an Invitrogen ATP detection kit (Invitrogen, A22066) according to their protocol. N=3 experiments, biological triplicates per genotype per experiment. Significance was calculated using a T-Test.

## Supplemental Information titles and legends

**Supplemental Figure 1: Related to Figure 1. A.** Comparison of gene and transcript structure of the *sloth1* and *sloth2* open reading frames. A common primer pair is used to distinguish genomic from cDNA (transcript) template by PCR. Sequence of *sloth1-2* genomic and *sloth1-2* transcript region provided. **B.** DNA gel image of PCR fragments amplified from indicated template samples. Predicted spliced transcript containing both sloth1 and sloth2 open reading frames is amplified from cDNA generated from adult flies, 3^rd^ instar larvae, and S2R+ cells.

**Supplemental Figure 2: Related to Figure 2. A.** Extended gene structure of *sloth1* and *sloth2* and genetic reagents. **B.** Sequence analysis of KO, dKO, and Gal4-KI alleles. **C.** (Left) Diagram of HDR knock-in of Gal4 into the *sloth1-sloth2* locus. (Right) DNA gel confirming Gal4 knock-in by PCR primers that flank the homology arms. Expected DNA fragment size in parenthesis.

**Supplemental Figure 3. Related to Figure 4.** Traces of electrical recordings from 3^rd^ instar larval NMJ in *dKO*, and *dKO+genomic rescue* animals. Graph on right is a quantification of the excitatory junction potential (EJP) for indicated genotypes. Significance was calculated with a T-Test compared to the *yw* control sample. Error bars show mean with SD. N ≥ 5 larvae per genotype.

**Supplemental Figure 4. Related to Figure 5.** Confocal microscopy images of 3^rd^ instar larval NMJ at muscle 6/7 segment A2. Antibodies or fluorescent proteins (green) mark synaptic components and anti-HRP (red) marks neurons. Comparison of wild-type to dKO. Graph shows quantification of synaptic bouton number by anti-Dlg1 staining. Significance of dKO bouton number was calculated with a T-test compared to WT. Error bars show mean with SD. N ≥ 7 NMJs (each from a different animal).

**Supplemental Figure 5. Related to Figure 5. A-C.** Transmission electron microscopy (TEM) images of sectioned adult eye photoreceptors from indicated genetic backgrounds with accompanying quantification of photoreceptor number and aberrant photoreceptors. Scalebar is 2µm. Filled red arrows indicate dead or dying photoreceptors. Open red arrows indicate unhealthy photoreceptors. Error bars show mean with SD. **A.** Animals were 4 weeks old and raised in a 12hr light/dark cycle. **B.** Animals were 1-3 days old and raised in a 12hr light/dark cycle. **C.** Animals were 4 weeks old and raised in the dark.

**Supplemental Figure 6. Related to Figure 5.** Confocal microscopy of adult eye photoreceptors stained with phalloidin (green) and anti-Rh1 (red). Animals were 4 weeks old and raised in the dark. Arrows indicate photoreceptors with higher levels of Rh1.

**Supplemental Figure 7. Related to Figure 6.** Confocal microscopy of 3^rd^ instar larval brain with antibody staining. Anti-Sloth1 or Anti-Sloth2 (green), mitochondria labeled with anti-ATP5alpha (red), and nuclei labeled with DAPI (blue). Wild-type (*yw*) or *sloth1/2* KO. **A.** Zoom out of entire brain showing region imaged in panels B and C. Scale bar 100µm. **B.** Results using two independent anti-Sloth1 antibodies (#1 and #2). Scale bar 20µm. **C.** Results using two independent anti-Sloth2 antibodies (#1 and #2). Scale bar 20µm.

**Supplemental Figure 8. Related to Figure 6.** SDS-PAGE and western blotting using anti-Sloth1 and anti-Sloth2 antibodies of cell and mitochondrial lysates. Two independent (#1 and #2) anti-Sloth1 and Anti-Sloth2 antibodies were tested. Arrowheads indicated expected band, asterisks indicate unrelated band(s). Tricine gels were used. **A.** S2R+ whole cell lysates isolated from indicated genotypes. Rhodamine-Actin used as loading control. **B.** S2R+ mitochondrial lysates isolated from indicated genotypes. Anti-ATP5alpha used as loading control. Mitochondrial control = ATP5alpha, cytoplasmic control = alpha-tubulin. **C.** S2R+ fractions isolated from wild-type S2R+ cells. WCL = Whole Cell Lysate, cyto. = cytoplasmic lysate, mito. = mitochondrial lysate. Blots were stripped and reprobed after detection of each antigen. **D.** Mitochondrial lysates isolated from 3^rd^ instar larvae or adult thorax mitochondrial isolation of indicated genotypes. “da>” indicates da-Gal4 crossed with attP40 (wild-type), RNAi (UAS-shRNA- sloth1/2), OE (UAS-sloth1/2 transcript).

**Supplemental Figure 9. Related to Figure 7. A.** Sequence analysis of single KO S2R+ clones for *sloth1* (clone 2F8) and *sloth2* (clone 3A7). sgRNA and PAM site indicated by grey boxes. **B.** PCR genotyping of four independently derived single cell dKO S2R+ clones. **C-D.** Seahorse mitochondrial stress test quantification of **C.** ATP production and **D.** Proton leak. Significance of KO lines was calculated with a T-test compared to S2R+. Error bars show mean with SD. ** P≤0.01, *** P≤0.001, **** P≤0.0001. N=6 for each genotype. **E.** Confocal images of 3^rd^ instar larval ventral nerve cord (VNC), axon bundles, and neuromuscular junction (NMJ). *MN-Gal4 UAS-mitoGFP* (*MN>mitoGFP*) (GFP) expresses mitochondrial-localized GFP in motor neurons. Neurons are stained with anti-HRP (magenta).

**Supplemental Figure 10. Related to Figure 7. A-B.** TEM images of sectioned adult photoreceptors. **A.** Adult flies are 4 weeks old and raised on a 12hr light/dark cycle. Mitochondria are indicated with red dots. **B.** Adult flies are 3 days old and raised in a 12hr light/dark cycle.

**Supplemental File 1.** Genomic sequence of *sloth1-sloth2* homologs in *D. melanogaster*, *S. urceolata*, *P. marinus*, and *C. intestinalis*

**Supplemental File 2.** Oligo and dsDNA sequences

**Supplemental File 3.** Gateway cloning plasmid list

**Supplemental File 4.** Raw gel and western images

---

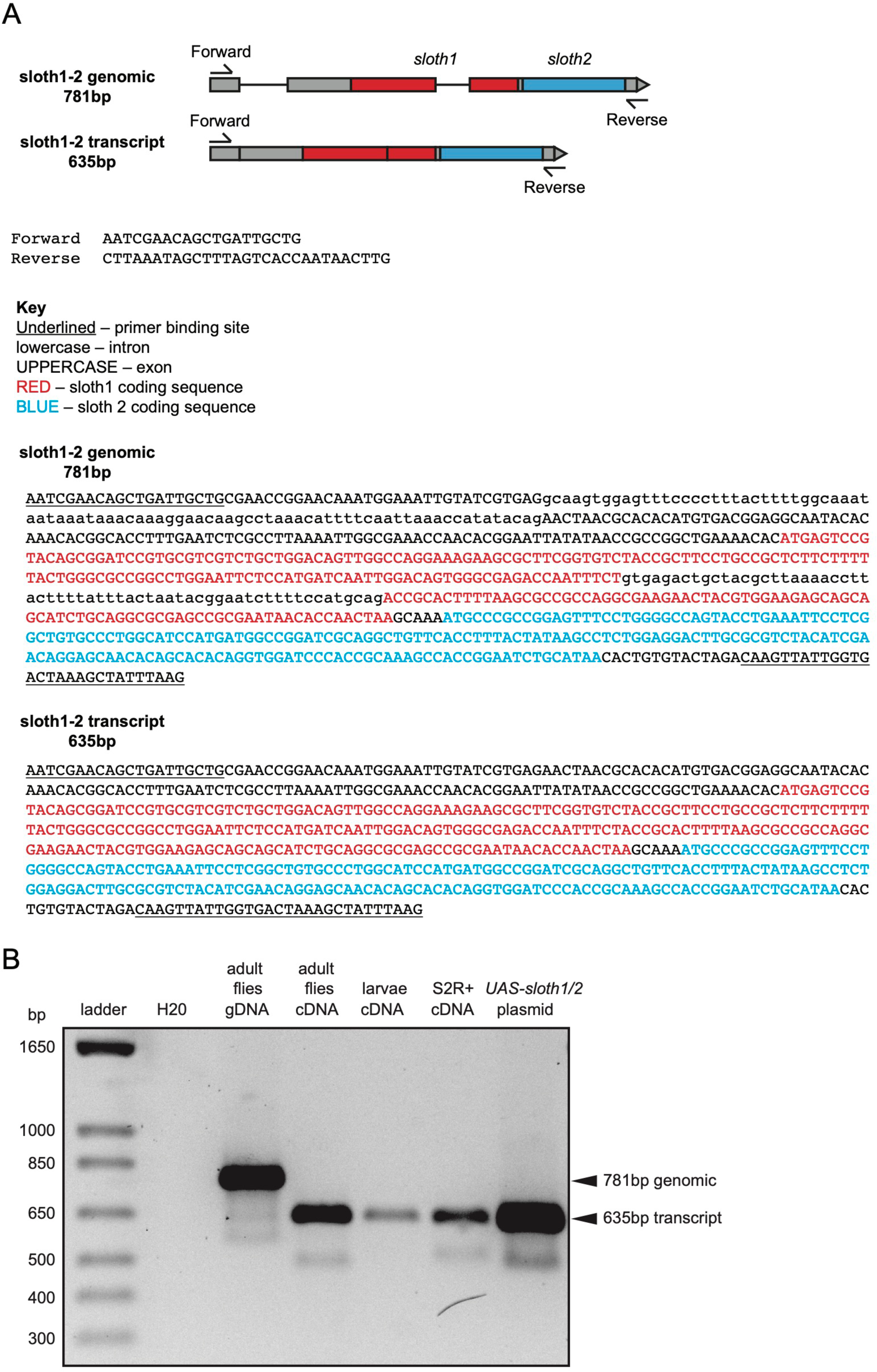

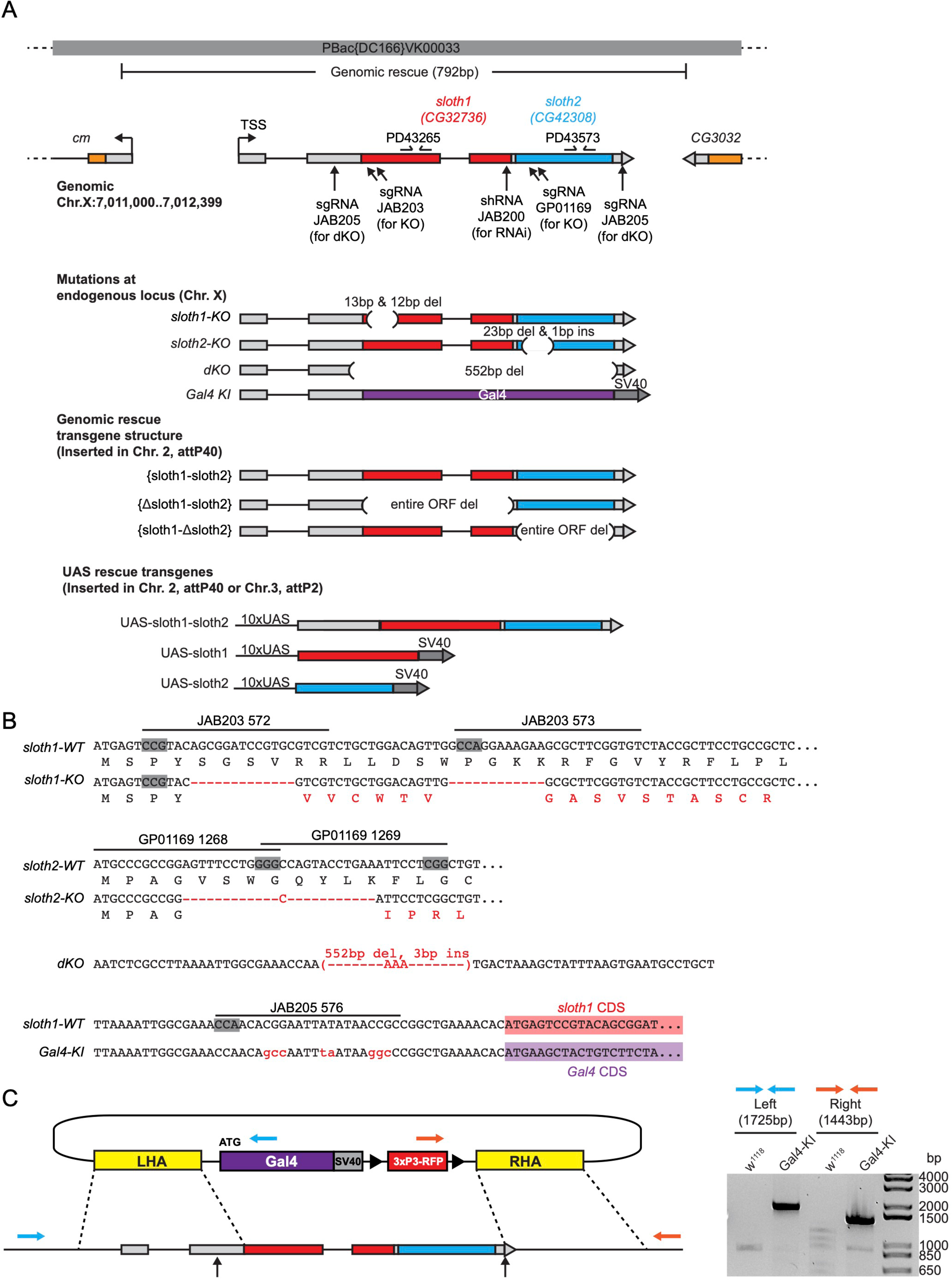

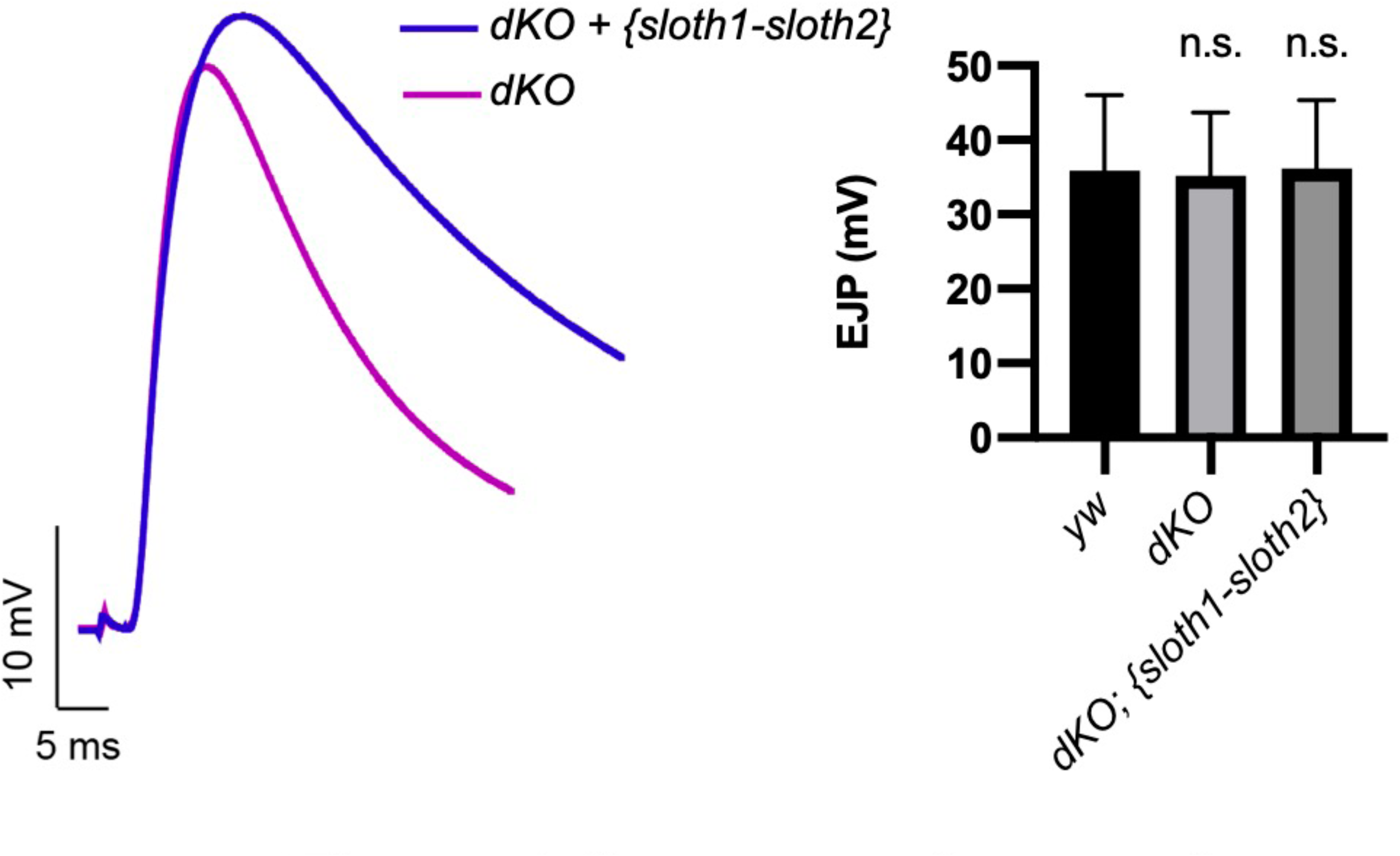

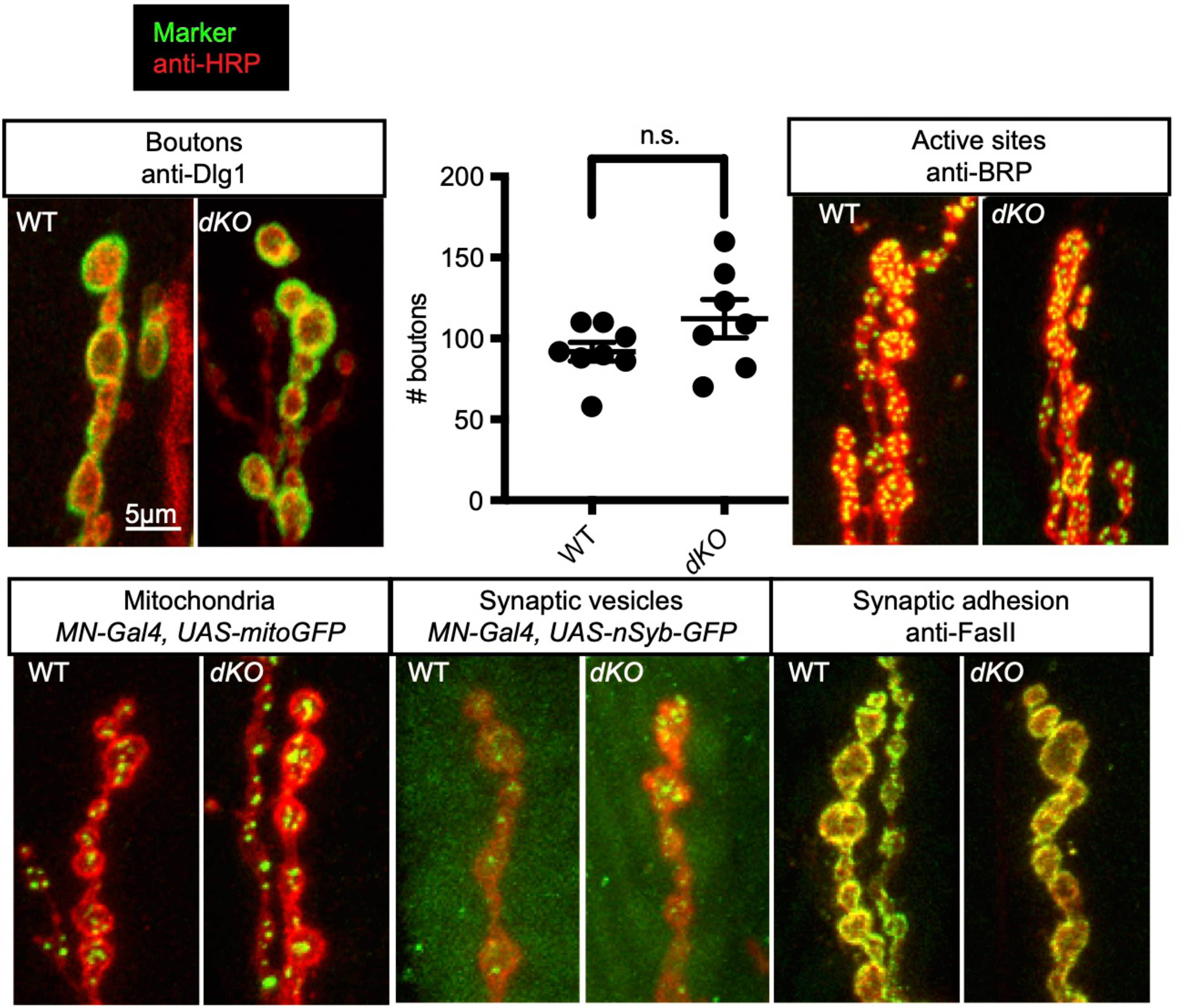

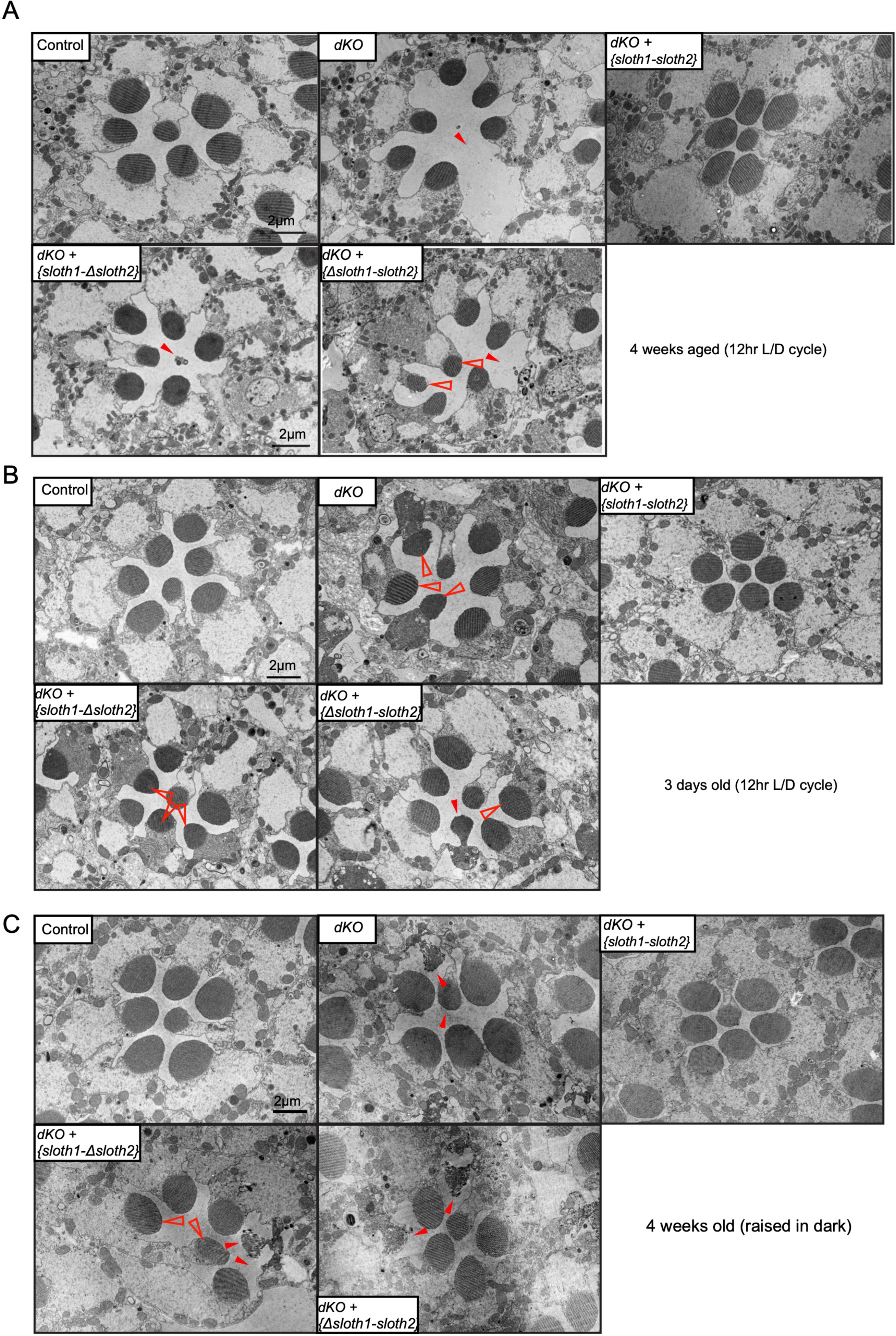

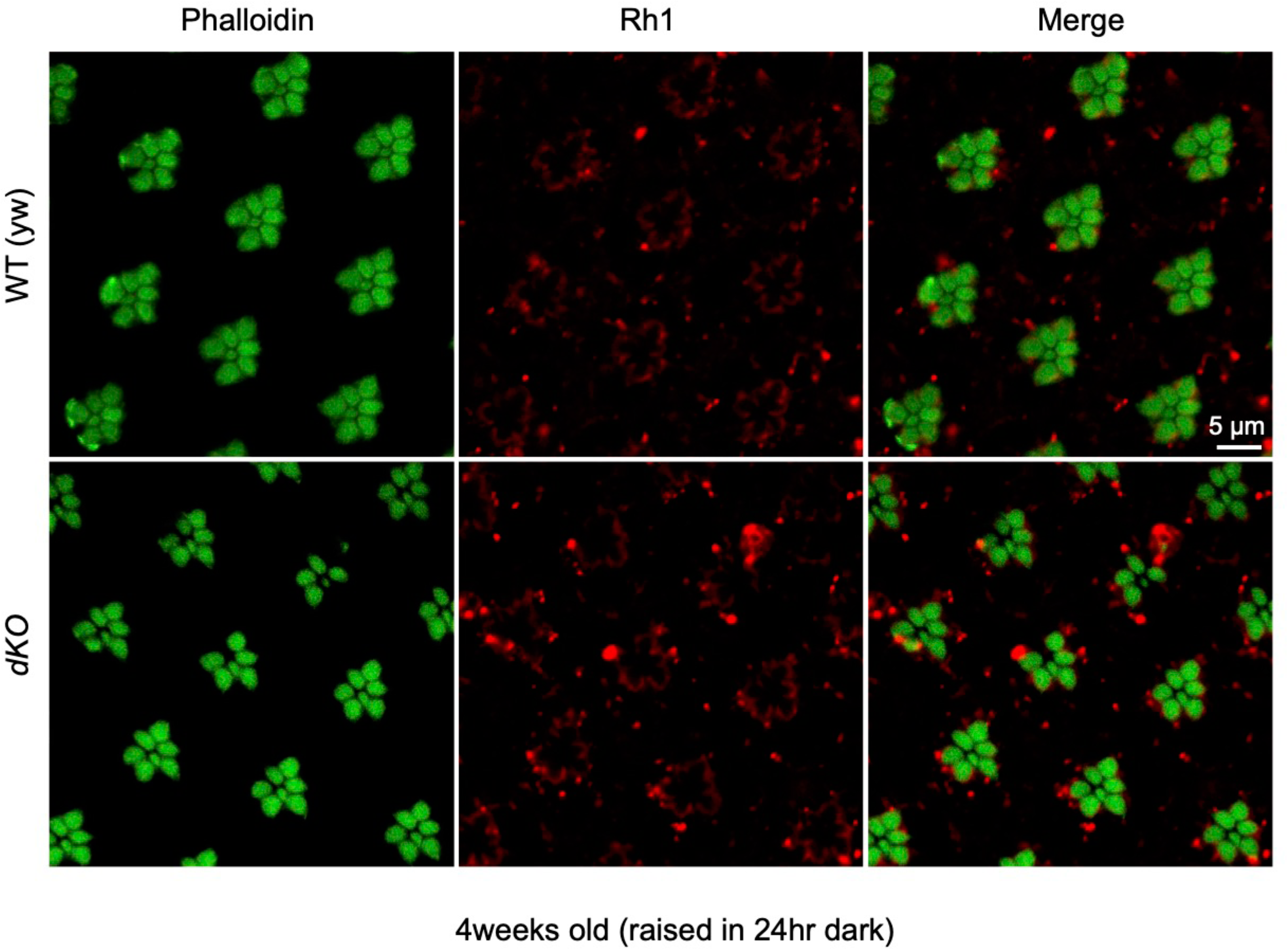

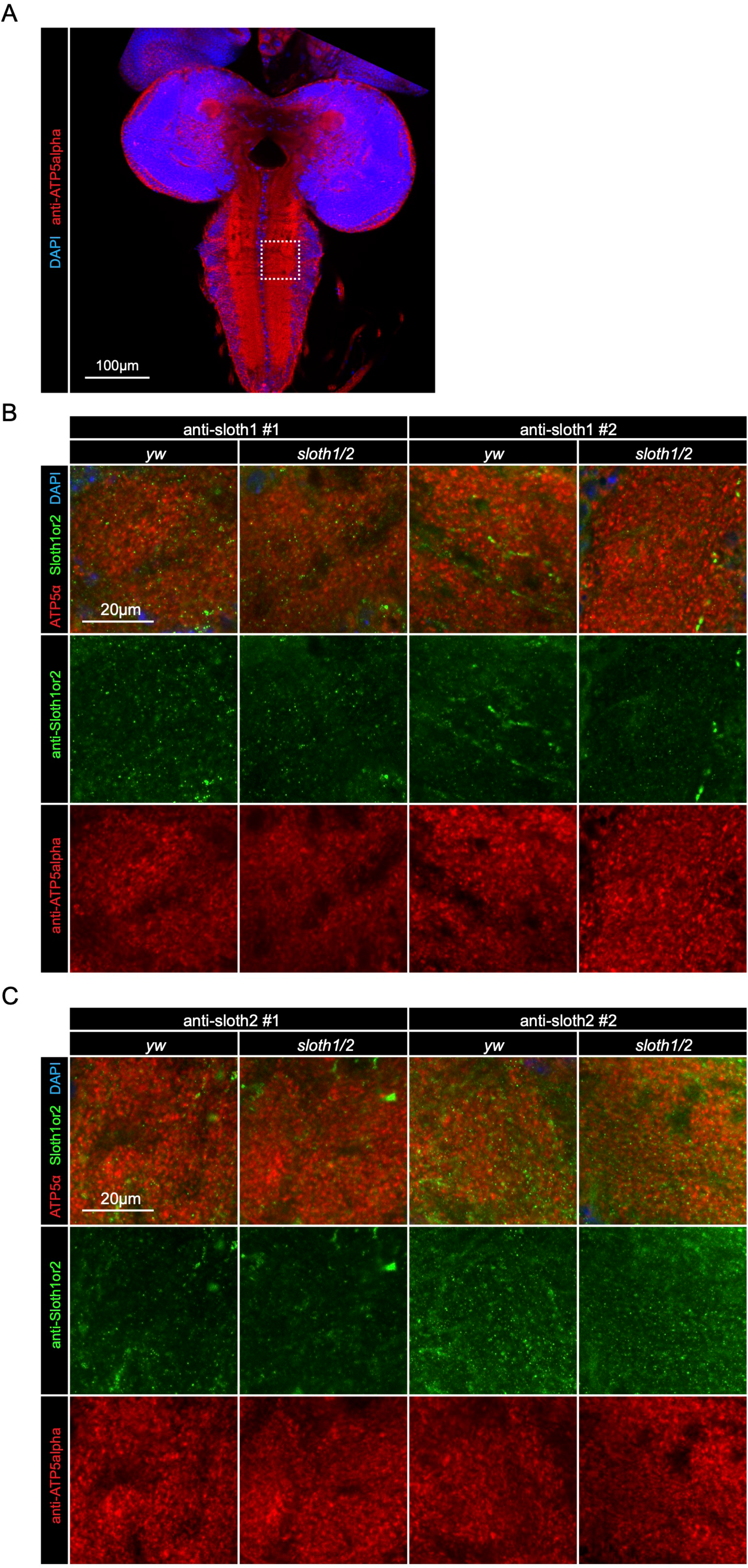

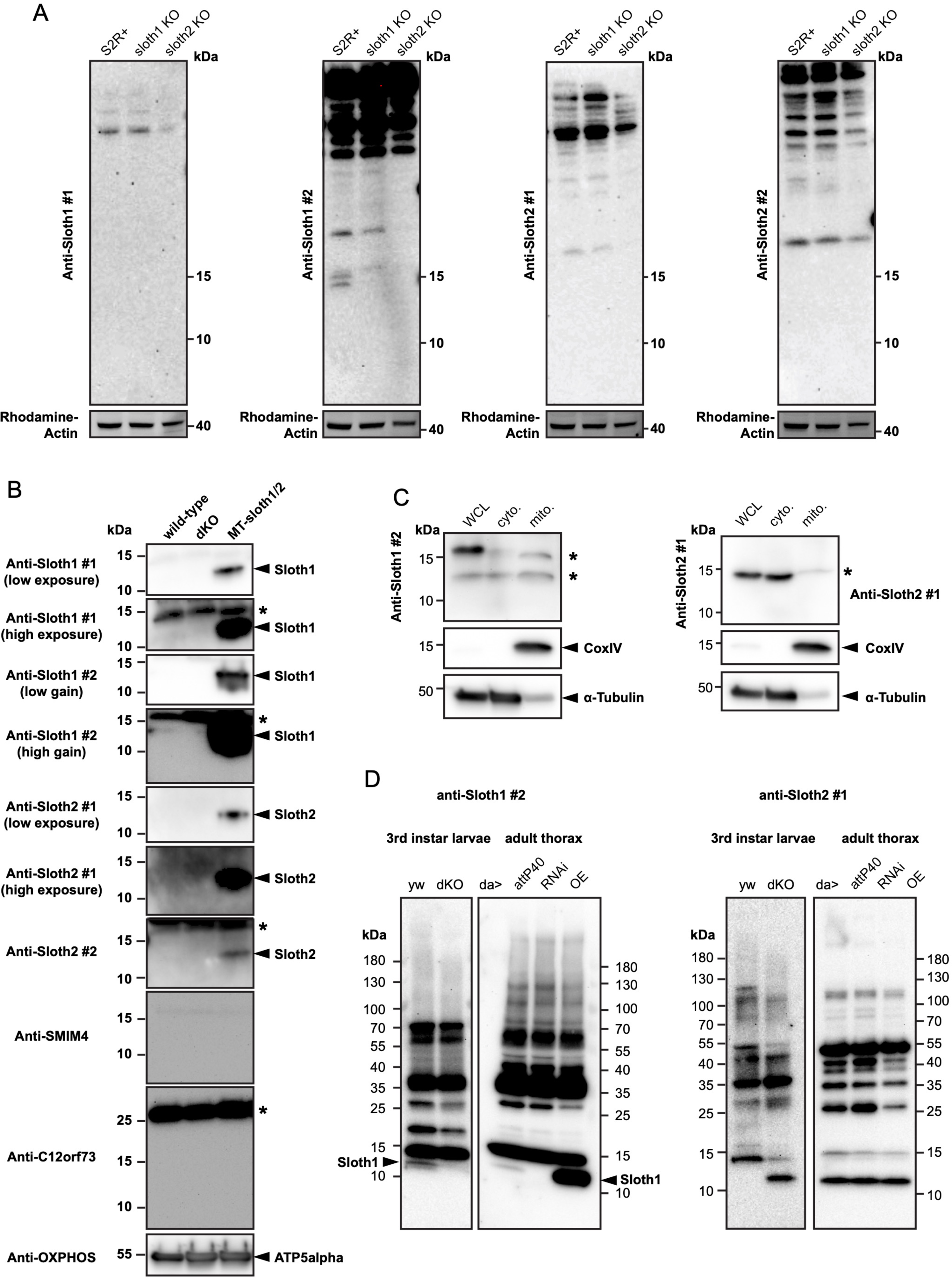

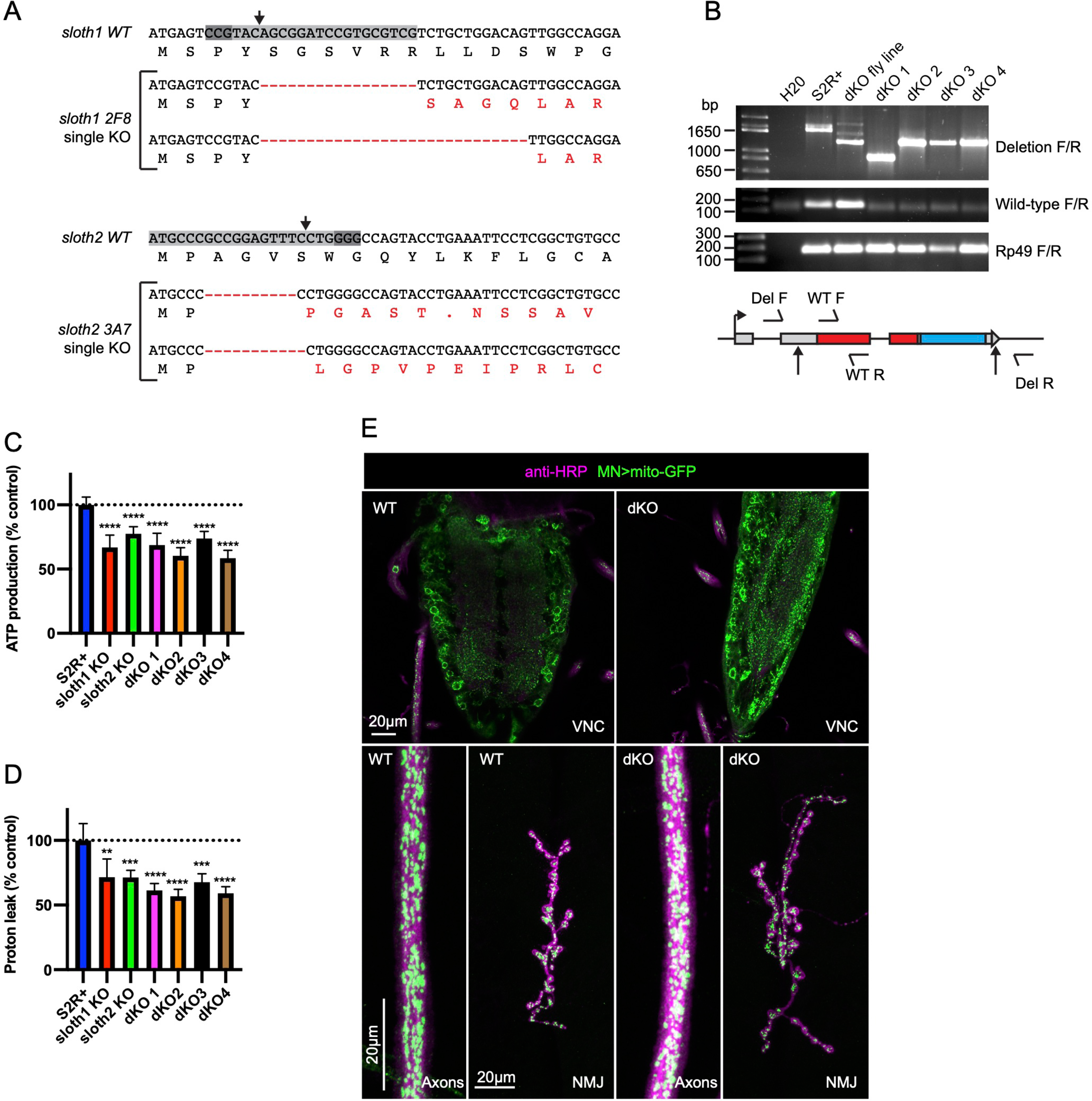

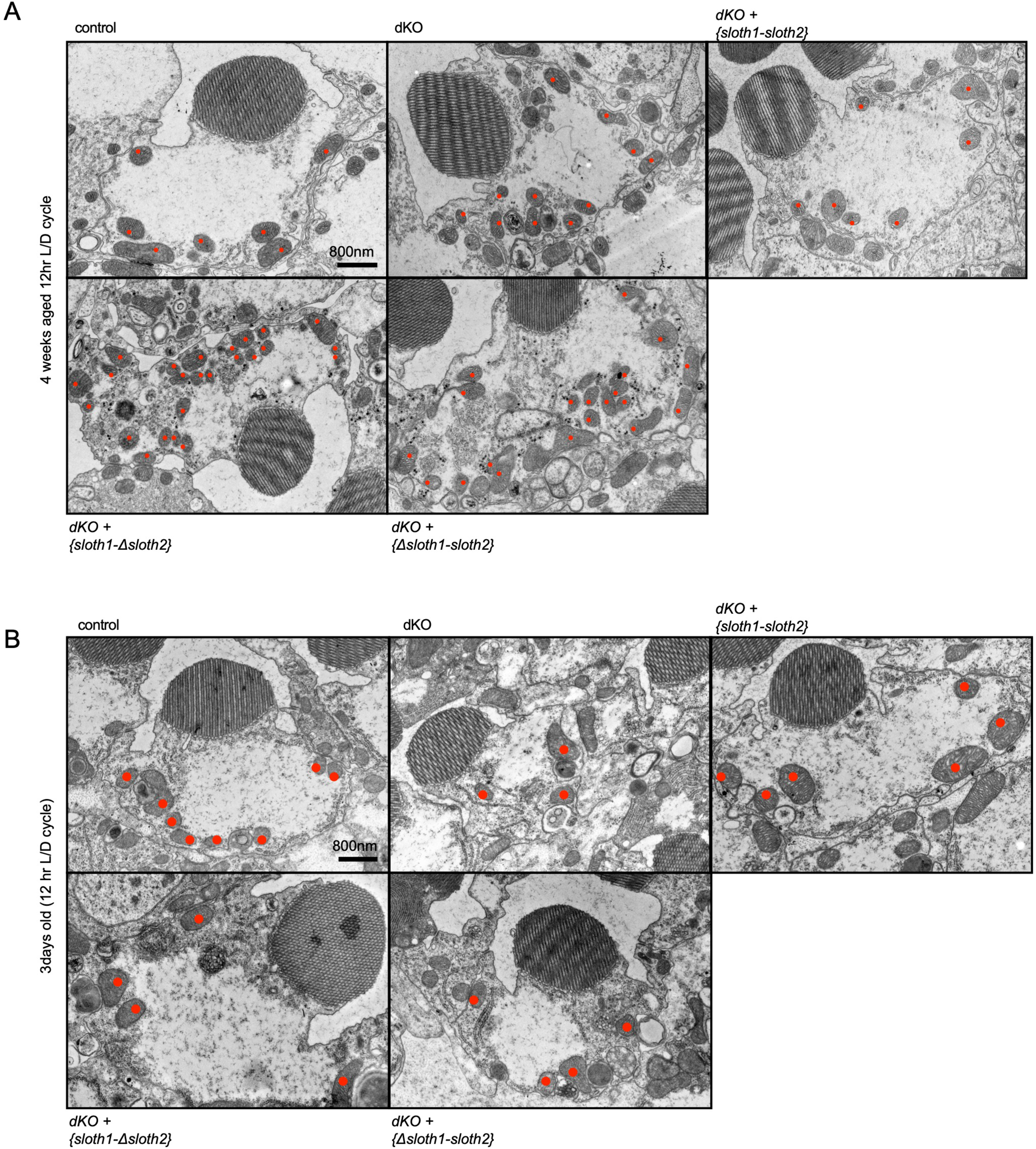

## References

Alloway, P. G., L. Howard and P. J. Dolph, 2000 The formation of stable rhodopsin-arrestin complexes induces apoptosis and photoreceptor cell degeneration. Neuron 28: 129–138.

Anderson, D. M., K. M. Anderson, C. L. Chang, C. A. Makarewich, B. R. Nelson et al., 2015 A micropeptide encoded by a putative long noncoding RNA regulates muscle performance. Cell 160: 595–606.

Antonicka, H., Z. Y. Lin, A. Janer, M. J. Aaltonen, W. Weraarpachai et al., 2020 A High-Density Human Mitochondrial Proximity Interaction Network. Cell Metab 32: 479–497 e479.

Basrai, M. A., P. Hieter and J. D. Boeke, 1997 Small open reading frames: beautiful needles in the haystack. Genome Res 7: 768–771.

Bergendahl, L. T., L. Gerasimavicius, J. Miles, L. Macdonald, J. N. Wells et al., 2019 The role of protein complexes in human genetic disease. Protein Sci 28: 1400–1411.

Bi, P., A. Ramirez-Martinez, H. Li, J. Cannavino, J. R. McAnally et al., 2017 Control of muscle formation by the fusogenic micropeptide myomixer. Science 356: 323–327.

Blumenthal, T., 2004 Operons in eukaryotes. Brief Funct Genomic Proteomic 3: 199–211.

Bosch, J. A., N. H. Tran and I. K. Hariharan, 2015 CoinFLP: a system for efficient mosaic screening and for visualizing clonal boundaries in Drosophila. Development 142: 597–606.

Brand, A. H., and N. Perrimon, 1993 Targeted gene expression as a means of altering cell fates and generating dominant phenotypes. Development 118: 401–415.

Brent, J. R., K. M. Werner and B. D. McCabe, 2009 Drosophila larval NMJ dissection. J Vis Exp.

Busch, J. D., M. Cipullo, I. Atanassov, A. Bratic, E. Silva Ramos et al., 2019 MitoRibo- Tag Mice Provide a Tool for In Vivo Studies of Mitoribosome Composition. Cell Rep 29: 1728–1738 e1729.

Calvo, S. E., K. R. Clauser and V. K. Mootha, 2016 MitoCarta2.0: an updated inventory of mammalian mitochondrial proteins. Nucleic Acids Res 44: D1251–1257.

Casson, S. A., P. M. Chilley, J. F. Topping, I. M. Evans, M. A. Souter et al., 2002 The POLARIS gene of Arabidopsis encodes a predicted peptide required for correct root growth and leaf vascular patterning. Plant Cell 14: 1705–1721.

Cavener, D. R., 1987 Comparison of the consensus sequence flanking translational start sites in Drosophila and vertebrates. Nucleic Acids Res 15: 1353–1361.

Chen, C. L., Y. Hu, N. D. Udeshi, T. Y. Lau, F. Wirtz-Peitz et al., 2015 Proteomic mapping in live Drosophila tissues using an engineered ascorbate peroxidase. Proc Natl Acad Sci U S A 112: 12093–12098.

Chen, J., A. D. Brunner, J. Z. Cogan, J. K. Nunez, A. P. Fields et al., 2020 Pervasive functional translation of noncanonical human open reading frames. Science 367: 1140–1146.

Chng, S. C., L. Ho, J. Tian and B. Reversade, 2013 ELABELA: a hormone essential for heart development signals via the apelin receptor. Dev Cell 27: 672–680.

Chugunova, A., E. Loseva, P. Mazin, A. Mitina, T. Navalayeu et al., 2019 LINC00116 codes for a mitochondrial peptide linking respiration and lipid metabolism. Proc Natl Acad Sci U S A 116: 4940–4945.

Couso, J. P., and P. Patraquim, 2017 Classification and function of small open reading frames. Nat Rev Mol Cell Biol 18: 575–589.

Crosby, M. A., L. S. Gramates, G. Dos Santos, B. B. Matthews, S. E. St Pierre et al., 2015 Gene Model Annotations for Drosophila melanogaster: The Rule-Benders. G3 (Bethesda) 5: 1737-1749.

Dennerlein, S., S. Poerschke, S. Oeljeklaus, C. Wang, R. Richter-Dennerlein et al., 2021 Defining the interactome of the human mitochondrial ribosome identifies SMIM4 and TMEM223 as respiratory chain assembly factors. Elife 10.

Fricker, L. D., 2005 Neuropeptide-processing enzymes: applications for drug discovery. AAPS J 7: E449–455.

Fukasawa, Y., J. Tsuji, S. C. Fu, K. Tomii, P. Horton et al., 2015 MitoFates: improved prediction of mitochondrial targeting sequences and their cleavage sites. Mol Cell Proteomics 14: 1113–1126.

Galindo, M. I., J. I. Pueyo, S. Fouix, S. A. Bishop and J. P. Couso, 2007 Peptides encoded by short ORFs control development and define a new eukaryotic gene family. PLoS Biol 5: e106.

Garcia, C. J., J. Khajeh, E. Coulanges, E. I. Chen and E. Owusu-Ansah, 2017 Regulation of Mitochondrial Complex I Biogenesis in Drosophila Flight Muscles. Cell Rep 20: 264–278.

Golpich, M., E. Amini, Z. Mohamed, R. Azman Ali, N. Mohamed Ibrahim et al., 2017 Mitochondrial Dysfunction and Biogenesis in Neurodegenerative diseases: Pathogenesis and Treatment. CNS Neurosci Ther 23: 5–22.

Gratz, S. J., F. P. Ukken, C. D. Rubinstein, G. Thiede, L. K. Donohue et al., 2014 Highly specific and efficient CRISPR/Cas9-catalyzed homology-directed repair in Drosophila. Genetics 196: 961–971.

Guo, X., A. Chavez, A. Tung, Y. Chan, C. Kaas et al., 2018 High-throughput creation and functional profiling of DNA sequence variant libraries using CRISPR-Cas9 in yeast. Nat Biotechnol 36: 540–546.

Hardie, R. C., and P. Raghu, 2001 Visual transduction in Drosophila. Nature 413: 186–193.

Hoskins, R. A., J. M. Landolin, J. B. Brown, J. E. Sandler, H. Takahashi et al., 2011 Genome-wide analysis of promoter architecture in Drosophila melanogaster. Genome Res 21: 182–192.

Hsu, P. Y., and P. N. Benfey, 2018 Small but Mighty: Functional Peptides Encoded by Small ORFs in Plants. Proteomics 18: e1700038.

Jaiswal, M., N. A. Haelterman, H. Sandoval, B. Xiong, T. Donti et al., 2015 Impaired Mitochondrial Energy Production Causes Light-Induced Photoreceptor Degeneration Independent of Oxidative Stress. PLoS Biol 13: e1002197.

Kann, O., and R. Kovacs, 2007 Mitochondria and neuronal activity. Am J Physiol Cell Physiol 292: C641–657.

Karginov, T. A., D. P. H. Pastor, B. L. Semler and C. M. Gomez, 2017 Mammalian Polycistronic mRNAs and Disease. Trends Genet 33: 129–142.

Katsir, L., K. A. Davies, D. C. Bergmann and T. Laux, 2011 Peptide signaling in plant development. Curr Biol 21: R356–364.

Kumar, S., Y. Yoshida and M. Noda, 1993 Cloning of a cDNA which encodes a novel ubiquitin-like protein. Biochem Biophys Res Commun 195: 393–399.

Liu, X., K. Salokas, F. Tamene, Y. Jiu, R. G. Weldatsadik et al., 2018 An AP-MS- and BioID-compatible MAC-tag enables comprehensive mapping of protein interactions and subcellular localizations. Nat Commun 9: 1188.

Magny, E. G., J. I. Pueyo, F. M. Pearl, M. A. Cespedes, J. E. Niven et al., 2013 Conserved regulation of cardiac calcium uptake by peptides encoded in small open reading frames. Science 341: 1116–1120.

Makarewich, C. A., K. K. Baskin, A. Z. Munir, S. Bezprozvannaya, G. Sharma et al., 2018 MOXI Is a Mitochondrial Micropeptide That Enhances Fatty Acid beta- Oxidation. Cell Rep 23: 3701–3709.

Mudge, J. M., J. Ruiz-Orera, J. R. Prensner, M. A. Brunet, F. Calvet et al., 2022 Standardized annotation of translated open reading frames. Nat Biotechnol 40: 994–999.

Murari, A., S. K. Rhooms, N. S. Goparaju, M. Villanueva and E. Owusu-Ansah, 2020 An antibody toolbox to track complex I assembly defines AIF’s mitochondrial function. J Cell Biol 219.

Nelson, B. R., C. A. Makarewich, D. M. Anderson, B. R. Winders, C. D. Troupes et al., 2016 A peptide encoded by a transcript annotated as long noncoding RNA enhances SERCA activity in muscle. Science 351: 271–275.

Ni, J. Q., R. Zhou, B. Czech, L. P. Liu, L. Holderbaum et al., 2011 A genome-scale shRNA resource for transgenic RNAi in Drosophila. Nat Methods 8: 405–407.

Nickless, A., J. M. Bailis and Z. You, 2017 Control of gene expression through the nonsense-mediated RNA decay pathway. Cell Biosci 7: 26.

Papp, B., C. Pal and L. D. Hurst, 2003 Dosage sensitivity and the evolution of gene families in yeast. Nature 424: 194–197.

Pauli, A., M. L. Norris, E. Valen, G. L. Chew, J. A. Gagnon et al., 2014 Toddler: an embryonic signal that promotes cell movement via Apelin receptors. Science 343: 1248636.

Pearson, R. K., B. Anderson and J. E. Dixon, 1993 Molecular biology of the peptide hormone families. Endocrinol Metab Clin North Am 22: 753–774.

Pellegrino, M. W., A. M. Nargund and C. M. Haynes, 2013 Signaling the mitochondrial unfolded protein response. Biochim Biophys Acta 1833: 410–416.

Perkins, L. A., L. Holderbaum, R. Tao, Y. Hu, R. Sopko et al., 2015 The Transgenic RNAi Project at Harvard Medical School: Resources and Validation. Genetics 201: 843–852.

Plaza, S., G. Menschaert and F. Payre, 2017 In Search of Lost Small Peptides. Annu Rev Cell Dev Biol 33: 391–416.

Port, F., H. M. Chen, T. Lee and S. L. Bullock, 2014 Optimized CRISPR/Cas tools for efficient germline and somatic genome engineering in Drosophila. Proc Natl Acad Sci U S A 111: E2967–2976.

Prelich, G., 2012 Gene overexpression: uses, mechanisms, and interpretation. Genetics 190: 841–854.

Pueyo, J. I., E. G. Magny, C. J. Sampson, U. Amin, I. R. Evans et al., 2016 Hemotin, a Regulator of Phagocytosis Encoded by a Small ORF and Conserved across Metazoans. PLoS Biol 14: e1002395.

Richter, D. J., P. Fozouni, M. B. Eisen and N. King, 2018 Gene family innovation, conservation and loss on the animal stem lineage. Elife 7.

Saghatelian, A., and J. P. Couso, 2015 Discovery and characterization of smORF- encoded bioactive polypeptides. Nat Chem Biol 11: 909–916.

Sandoval, H., C. K. Yao, K. Chen, M. Jaiswal, T. Donti et al., 2014 Mitochondrial fusion but not fission regulates larval growth and synaptic development through steroid hormone production. Elife 3.

Sardiello, M., F. Licciulli, D. Catalano, M. Attimonelli and C. Caggese, 2003 MitoDrome: a database of Drosophila melanogaster nuclear genes encoding proteins targeted to the mitochondrion. Nucleic Acids Res 31: 322–324.

Savojardo, C., N. Bruciaferri, G. Tartari, P. L. Martelli and R. Casadio, 2020 DeepMito: accurate prediction of protein sub-mitochondrial localization using convolutional neural networks. Bioinformatics 36: 56–64.

Snyder, S. H., and R. B. Innis, 1979 Peptide neurotransmitters. Annu Rev Biochem 48: 755–782.

Sopko, R., D. Huang, N. Preston, G. Chua, B. Papp et al., 2006 Mapping pathways and phenotypes by systematic gene overexpression. Mol Cell 21: 319–330.

Stapleton, M., J. Carlson, P. Brokstein, C. Yu, M. Champe et al., 2002 A Drosophila full- length cDNA resource. Genome Biol 3: RESEARCH0080.

Stein, C. S., P. Jadiya, X. Zhang, J. M. McLendon, G. M. Abouassaly et al., 2018 Mitoregulin: A lncRNA-Encoded Microprotein that Supports Mitochondrial Supercomplexes and Respiratory Efficiency. Cell Rep 23: 3710–3720 e3718.

Suzuki, K., T. Hashimoto and E. Otaka, 1990 Yeast ribosomal proteins: XI. Molecular analysis of two genes encoding YL41, an extremely small and basic ribosomal protein, from Saccharomyces cerevisiae. Curr Genet 17: 185–190.

Szklarczyk, R., M. A. Huynen and B. Snel, 2008 Complex fate of paralogs. BMC Evol Biol 8: 337.

Taylor, J. S., and J. Raes, 2004 Duplication and divergence: the evolution of new genes and old ideas. Annu Rev Genet 38: 615–643.

Thompson, S. R., 2012 So you want to know if your message has an IRES? Wiley Interdiscip Rev RNA 3: 697–705.

Thul, P. J., L. Akesson, M. Wiking, D. Mahdessian, A. Geladaki et al., 2017 A subcellular map of the human proteome. Science 356.

Trevisan, T., D. Pendin, A. Montagna, S. Bova, A. M. Ghelli et al., 2018 Manipulation of Mitochondria Dynamics Reveals Separate Roles for Form and Function in Mitochondria Distribution. Cell Rep 23: 1742–1753.

Ugur, B., H. Bao, M. Stawarski, L. R. Duraine, Z. Zuo et al., 2017 The Krebs Cycle Enzyme Isocitrate Dehydrogenase 3A Couples Mitochondrial Metabolism to Synaptic Transmission. Cell Rep 21: 3794–3806.

Usui, S., L. Yu and C. A. Yu, 1990 The small molecular mass ubiquinone-binding protein (QPc-9.5 kDa) in mitochondrial ubiquinol-cytochrome c reductase: isolation, ubiquinone-binding domain, and immunoinhibition. Biochemistry 29: 4618–4626.

Van Der Kelen, K., R. Beyaert, D. Inze and L. De Veylder, 2009 Translational control of eukaryotic gene expression. Crit Rev Biochem Mol Biol 44: 143–168.

Veitia, R. A., S. Bottani and J. A. Birchler, 2008 Cellular reactions to gene dosage imbalance: genomic, transcriptomic and proteomic effects. Trends Genet 24: 390–397.

Verstreken, P., C. V. Ly, K. J. Venken, T. W. Koh, Y. Zhou et al., 2005 Synaptic mitochondria are critical for mobilization of reserve pool vesicles at Drosophila neuromuscular junctions. Neuron 47: 365–378.

Viswanatha, R., Z. Li, Y. Hu and N. Perrimon, 2018 Pooled genome-wide CRISPR screening for basal and context-specific fitness gene essentiality in Drosophila cells. Elife 7.

Wang, J. W., E. S. Beck and B. D. McCabe, 2012 A modular toolset for recombination transgenesis and neurogenetic analysis of Drosophila. PLoS One 7: e42102.

Wu, C. F., and F. Wong, 1977 Frequency characteristics in the visual system of Drosophila: genetic dissection of electroretinogram components. J Gen Physiol 69: 705–724.

Xue, Z., M. Ren, M. Wu, J. Dai, Y. S. Rong et al., 2014 Efficient gene knock-out and knock-in with transgenic Cas9 in Drosophila. G3 (Bethesda) 4: 925–929.

Yamamoto, S., M. Jaiswal, W. L. Charng, T. Gambin, E. Karaca et al., 2014 A drosophila genetic resource of mutants to study mechanisms underlying human genetic diseases. Cell 159: 200–214.

Yanagawa, S., J. S. Lee and A. Ishimoto, 1998 Identification and characterization of a novel line of Drosophila Schneider S2 cells that respond to wingless signaling. J Biol Chem 273: 32353–32359.

Yang, L., and A. Veraksa, 2017 Single-Step Affinity Purification of ERK Signaling Complexes Using the Streptavidin-Binding Peptide (SBP) Tag. Methods Mol Biol 1487: 113–126.

Yeasmin, F., T. Yada and N. Akimitsu, 2018 Micropeptides Encoded in Transcripts Previously Identified as Long Noncoding RNAs: A New Chapter in Transcriptomics and Proteomics. Front Genet 9: 144.

Zhang, S., B. Reljic, C. Liang, B. Kerouanton, J. C. Francisco et al., 2020 Mitochondrial peptide BRAWNIN is essential for vertebrate respiratory complex III assembly. Nat Commun 11: 1312.

Zhou, R., I. Hotta, A. M. Denli, P. Hong, N. Perrimon et al., 2008 Comparative analysis of argonaute-dependent small RNA pathways in Drosophila. Mol Cell 32: 592–599.

