## Supplemental File 1 for "Two neuronal peptides encoded from a single transcript regulate mitochondrial complex III in *Drosophila*"

Genomic sequences of *sloth1* and *sloth2* homologs

**BOLD = Coding sequence**
Red = *sloth1* homolog
Blue = *sloth2* homolog

>Dmel_sloth1-sloth2(CG32736-CG42308)
AATCGAACAGCTGATTGCTGCGAACCGGAACAAATGGAAATTGTATCGTGAGgcaagtg
gagtttcccctttacttttggcaaataataaataaacaaaggaacaagcctaaacattt
tcaattaaaccatatacagAACTAACGCACACATGTGACGGAGGCAATACACAAACACG
GCACCTTTGAATCTCGCCTTAAAATTGGCGAAACCAACACGGAATTATATAACCGCCGG
CTGAAAACAC**ATGAGTCCGTACAGCGGATCCGTGCGTCGTCTGCTGGACAGTTGGCCAG**
**GAAAGAAGCGCTTCGGTGTCTACCGCTTCCTGCCGCTCTTCTTTTTACTGGCGCCGGC**
**CTGGAATTCTCCATGATCAATTGGACAGTGGGCGAGACCAATTTCT**gtgagactgctac
gcttaaaccttacttttatttactaatacggaatcttttccatgcag**ACCGCACTTTT**
**AAGCGCCGCCAGGCGAAGAACTACGTGGAAGAGCAGCAGCATCTGCAGGCGCGAGCCGC**
**GAATAACACCAACTAA**GCAAA**ATGCCCGCCGGAGTTTCCTGGGGCCAGTACCTGAAATT**
**CCTCGGCTGTGCCCTGGCATCCATGATGGCCGGATCGCAGGCTGTTCACCTTTACTATA**
**AGCCTCTGGAGGACTTGCGCGTCTACATCGAACAGGAGCAACACAGCACACAGGTGGAT**
**CCCACCGCAAAGCCACCGGAATCTGCATAA**CACTGTGTACTAGACAAGTTATTGGTGAC
TAAAGCTATTTAAG

>Choanoflagellate_Salpingoeca_urceolata_sloth1-
sloth2_comp15074_c0_seq2
TTCACTTTCGTTTTCTTACTGTTTCAACGTTGCGACTGTGCTCTTCGGCTTCACGTGTT
CTTGCACCATCTGCTGTGGCACCCATTCAGCGCAGAGTTCAGCGGTCCACGCAGTGGCA
GCGGGCCAGGACACCACTTCTGCTTGGGTACCTCTA**ATGCCGCGTTCGTTTCCGCAAAT**
**TGCGGCGCGTGTGGTGCCTGTGTCGTTTGCTCTTGGCGCGTTTATGGAATGGTTCATGC**
**TCAACGTTCAAATTGGCCACGAAACCTTTTATGACACTGCAGTGAGGCTGGAAGCAAAG**
**CGACGGTTTGAACAACAGCAAGAGGAGCAGCAAAAAGCTAGCAACGACCCTTCGTCCGA**
**CTCACCGCCGCCAGCAGCATCCTAA**GAGTTGTTTGCTTCCTGAAGTAGTTTTAGTTTGT
ACCTGTTGTTTTTCGTTAGTTTTTTTGAAGGTTCCTTCACGTCCAGCACC**ATGCCGTTT**
**GGTGTTTCCATGTCTCGGTACGTGGGTGTGGTCGCACTTACCCTCGGGTCCATGCTGGC**
**CGGTGCTTCCACCGTACACTACTTCTACCAGCCCGACCTGACTGTGCCCACCGAGCCTC**
**CTCCGGCGCCGGATTCCGTGTTGAAAAAGCCACGGATAGCCTTGGTGTCGCCACGGCAG**
**CGTGCGACGGGAGAAGCAGACGATGGAAAACAGTGA**CCGGTCTTATGCGTGATTGGTAT
TAAACACATGGTCGTGTTCAAGATGAGGTTGTTGGTTGCCAGTGCCGCGGAAAACCCGC
AACATGGGCGCTTGTCCCAATACGTTTTTGCTGTGGGGTGTTCGTTTTTCTTTTTCCGG
TTGGTTGTTTCATCCTCATTCGCCACGCAGCAGCAAAAAGCAACAAGTCAACTCGATTG

>Lamprey-Petromyzon_marinus_sloth1-sloth2
TTTCTGTCTGTGCCCGCGTGTCTCTGTGTCCACATGTCTGTCTGTCCATGTGTCAGGGG
GTGCAGCGGGCGAATGGGCG**ATGGTGTTCTTCAGCAGCGCTCTCGGGAGGATTCTCAGT**
**AAAGTTCCCGGAGAGAAGAGGCTGGGTGTCTATCGGTTCCTGCCCGTGTTCTTCGTGAT**
**TGGCGGTGCCATGGAGTGGATCATGATTAACATGAGAGTCGGCAGAGAGACCTTCT**GTG
GGTACCACGCAGGGCTTCATTATTTCTCACTGAAATATTTTCCGGGTGACCGGTAGACT

GGAGTTGGTTGCACATGATTAGTATCCACGGCCTGGTAGCCCTGAACAGCGCCTACACT
GGAATCGGGACTCGCATGCCACGCGTTTGACTCTTCGTTTGACCCTTCGTTTGACCCCG
GCGTCCCATTATTTACCTCTGACACCGCATGCTCACCATCGAGTGCGACTAACCGCACG
CGACGGCGCGCTGTTTCTTTCAG**ACGACGTCTACAGACGCAAGCAGTCGGAGCGCCGTT**
**ACCAGCAGCGCCTCGCCGAGACCTCGCAGTCCAGCGGTTCCAACTAA**GAGTCTCGCCTT
TCTCGAACAGACGATCGACTCGGTCACCACCCCACACGTCACTCCGTCTCCTCCCCCCC
TTCCCGCCGTTGTTGCTGCCGCCGCTGCCACCACAACCGACTTGCGCTGCTTGCGTAGA
AGCTACGGGCGCAAAGAACTGACGGCTCGCACTGGGCCGTGCGTGAGACTTTCGGAGCG
AGGTTGTTGACA**ATGCCGGCGGGCGTGACGTGGCCGCGCTATCTCAAGATGCTGACCGC**
**GAGTCTCCTGTCAATGCTGGCAGGAGCGGAGGTGGTTCACCGCTACTACCGGCCAGACC**
**TG**GTACGTGGACTTTTTTTCTTTCGTTCTCAGGAGTCCGGCTCGGGGATATAAAATGTT
CACGTTATAAGCCATTTCATTGAGCTATCATATGTGATAACCAGGTCGCTTCTGAAAAA
GAGCTAAATTACTCATTGGGCCTTACCTAGTAAAAAAAAATCCCACTGAGTGTTTTCCG
GGTCTCTGGTTAAACCCAAGAAGGTGACTCGCAGTAGCCGCAACCATAGCGAAGGAGGT
ATACTTGATGTGGTGTGTTGGGTGCAGAAATACAGGACCCCAAGAGACGCTGCTACCCG
TAGTGTATCTGTGTGGATATCCGGTGTTAATTGCCATGTAAGAGTGGGTAAGAGGATAT
TTCGATAGTACCACCCCAACAGGGATAAAGAGGGGTTTCCACCGCATTGCTGTTGTTCA
CTGTTGCGGTTTCCCTCCCACACAG**AGCATCCCTGAGGTTCCGCCAGCGCCGGGGCAAC**
**TGCAGACGCGGCTGTTGGGCATCGAGGGCACAACGGGGACACCACTCAGTGGCACCAGG**
**GCTGCGGAGGAGG**AACGCAGCCATCCCTCGTGACGGCGTCCACTCCCTCAACCTCGAGC
ACGTGCACGTGCACGAGTTAACGCACACACGAACATGCACAGGAGGCACAGCACATGCA
CAGAATGTTATACCTCCTTCACGATGGTGAATCAAAAACGATAAGACTTTTTATTTTAC

>seasquirt_XM_018812254.2_sloth1-sloth2
TTCAAAACAGAACAGTTATCAAATGTATTATGTAAAAATGCAGTTGAGTATATGAGTAA
GCCAGTAGTACATAATATAAACCATACCCTCGGTCTGGAGCCACAAATACTTAAAACAA
ATACGGCTAATACTTTTTGTAATATTCTAGTAACAAAACCTGATTTTTAAACATATTTG
GCCCATTTTAGAGTTGTAAAGTATGAATTGTTTCTAGT**ATGACGTTTATTGGTCGACTG**
**GTCCAGACATTTCTTTACTACTACCCAATAAAAAGACAAAGCCCATACAAATTCGTTCC**
**ACTGTTTTTTGCCATTGGAGCGTCTGTGGAGTGGGTTATGATAAAAGTTCCGGCTGCAG**
**GACGAGGTGAAACATTTTACGACGTTTGGAGAAGAAATAGATCAGAAAAAGAATACAAG**
**CAGAGAATAATTGAAGAGAAATTTCAAGAAGCAATTAAAGCAAAAGAAAACTGTGAAAA**
**TTAA**TAAGCATATATTTGGCTTGTCTTAAACTGCATTAAACACTTAATTTAAATAAATT
ACCTTTGAAAAAATCAATAATTTACTTTTATTATAAGTTTAAACAGTTTTTTTAGCTTG
AACTTGCGTAAAGAAATTTAGGCCTAAAATTAAAAATCACCCAAAAACACTTTCTGTTC
ATTTAATAAGCAAAACCTTTTGTTTGATTTATTTTCCAACTGTATAATTTTGCATACCC
ACCACATC**ATGCCTTATGGTGTTTCTTGGCCATTCTACCTGAAAACAGTATCTTCTTCA**
**CTCATAGCAATGTTCCTGGGCTCACACAGTGTTCATATGTGGTACAGACCTGATCTATC**
**CATACCTGAGATCCCACCTAAAAAAGGGGAGCTTCACACAAAACTTTATACAACAAAAT**
**CAGAAAATTAA**ACGAATTCATTACTTTTGTTAATGTTTTTTTGGTAACCTTAATCCAGT
GTGCAGTTGTACTATACGCTTATTTTTTTTTTGGTAGCTTTGTTTCAGCTAGTTACTTG
TTTTCTATCAGGTATACTGGTAATGTTTTGGTTTACATTTATTTATGAAGAAGATAAGT
TTCCTTCTGCTAAGTAAAAGTTGGCATTTTAAATGTAATTCACTTTAAAAACCCATATT
TCAGTTTCATTTCATAACGCTTTTTGTGTTTGATCAATTTTTGGCTGTGAACAAATTTT
GTGTTTGTTTGACTCAACCTAAAAACATCTCCTTACTTATTAGGTTGACTGTATAGGGC
AAAGTAGTTTTCAAACATTGTATAACTTTTCAAGATGGCCGACAACCTTAGTGAAGAAT
GGTGGCAAACGGCAGTTTCTGATGAAGAAGAAGGCGCAAGTGATGATGGTGAACGAAAA
GAAATGAAACGTAAACTGAACGAACCGACTTCAGGAATAGTAGTTTCAGAAAACGAGGA

ACCAGAAGTGAAAAAGAAAAAAAAGGCGGAACAGAAAAAGAATTACTGAAGCTAAGCTTC
CCGATCAAGGGGATTCACCCACGATGTTACGAGATTATCTCAAACTTCACTTCAGTAAA
TTATCCAAGCTCGAATTTGAGGATATTTCGCTAACAGAATCCAATTTCACAGCATGCAA
TATCGACAAAGAACATACTACCACGTCGTATTTTAAACAAATCGCCCCCAAGTGGCATC
GTTTAAGCACAGCTCACAGTCACAAGATGTCGCCTCTGATCATCGTGGTTTGTGGCAAC
GCACTTCGAGCGTCGAAATTTAACACAGAAGCAAAGACTTTTAAGGGCAAAGATGCAAG
GTCGATAAAGCTATTTGCGCGCCACATGAAGATCGACGATCAAATCAAACTTCTGCGGG
AAAACGTCATTCATTTCGCCGTCGGCACACCGGAAAGAATCCGATCTCTTATCCTACAA
GATGCTCTCAGTTTAGAACACACTCGAGCGTTTGTCATCGATTGGAATTGGAGAGATGT
AAAACTAAAGCGTTTAATTGACATACGAGAGGCTCGTGCGTCGTTGATGAATTTGTTAA
AAGATTGCGTGATCCCAGCTTGTAAGAAACACCATGTAAAAATCGGGTTGTTTTGATTT
GAATTTGTGCAAAAAAATGAGGTTTTCTGACGTCATACAGGTTCAAAATTTGCTTGTGTG
CATGGCCCGTTTTTTTCAGTAAATGGTTTACGTTCATGCAATAAATTGCCATTTTAAGT
TAGTGTA
