## Supplemental File 2 for "Two neuronal peptides encoded from a single transcript regulate mitochondrial complex III in *Drosophila*"

| Name | Sequence | Description |
|---|---|---|
| JB2578_CG3242_transcript_F | AATCGAACAGCTGATTGCTG | PCR primers to amplify sloth1-2 genomic or transcript region |
| JB2579_CG3242_transcript_R | CTTAAATAGCTTTAGTCACCAATAACTTG | PCR primers to amplify sloth1-2 genomic or transcript region |
| JB749_MT-Rluc_backboneF | ATGACTTCGAAAGTTTATGATC | Cloning pMT-sloth1-Rluc |
| JB750_MT-Rluc_backboneR | GAATTCCCCTTTAGTTGCAC | Cloning pMT-sloth1-RLuc |
| JB751_CG3242_5'UTR_overlapMTRluc_F | gtgcaactaaaggggaattcAATCGAACAGCTGATTGC | Cloning pMT-sloth1-RLuc |
| JB752_CG3242_5'UTR_overlapMTRluc_R | tcataaactttcgaagtcatTTTGCTTAGTTGGTGTTATTC | Cloning pMT-sloth1-RLuc |
| JB753_CG32_SDM_ATG_TTG_F | ctgaaaacacTTGAGTCCGTACAGCGGATC | Cloning pMT-sloth1-RLuc derivatives by SDM |
| JB754_CG32_SDM_ATG_TTG_R | acggactcaaGTGTTTTCAGCCGGCGGTTATATAATTC | Cloning pMT-sloth1-RLuc derivatives by SDM |
| JB755_CG32_SDM_ATG_del_F | ctgaaaacacAGTCCGTACAGCGGATC | Cloning pMT-sloth1-RLuc derivatives by SDM |
| JB756_CG32_SDM_ATG_del_R | acggactGTGTTTTCAGCCGGCGGTTATATAATTC | Cloning pMT-sloth1-RLuc derivatives by SDM |
| JB757_CD32_SDM_kozak_GTGT_F | ctgaaagtgtATGAGTCCGTACAGCGGATC | Cloning pMT-sloth1-RLuc derivatives by SDM |
| JB758_CD32_SDM_kozak_GTGT_R | acggactcatACACTTTCAGCCGGCGGTTATATAATTC | Cloning pMT-sloth1-RLuc derivatives by SDM |
| JB759_CD32_SDM_kozak_CAAA_F | ctgaaacaaaATGAGTCCGTACAGCGGATC | Cloning pMT-sloth1-RLuc derivatives by SDM |
| JB760_CD32_SDM_kozak_CAAA_R | acggactcatTTTGTTTCAGCCGGCGGTTATATAATTC | Cloning pMT-sloth1-RLuc derivatives by SDM |
| JB773_CG42_SDM_kozak_GTGT_F | accaactaagGTGTATGACTTCGAAAGTTTATGATCCAG | Cloning pMT-sloth1-RLuc derivatives by SDM |
| JB774_CG42_SDM_kozak_GTGT_R | tcgaagtcatACACCTTAGTTGGTGTTATTCGC | Cloning pMT-sloth1-RLuc derivatives by SDM |
| JB567_CG32736_shRNA3_top | ctagcagtGCCGCGAATAACACCAACTAAtagttatattcaagcataTTAGTTGGTGTTATTCGCGGCgcg | Oligos annealed and ligated into pValium20 for shRNA expression |
| JB568_CG32736_shRNA3_bot | aattcgcGCCGCGAATAACACCAACTAAtatgcttgaatataactaTTAGTTGGTGTTATTCGCGGCactg | Oligos annealed and ligated into pValium20 for shRNA expression |
| JB572_CG32736gRNAdKO_F | TATATAGGAAAGATATCCGGGTGAACTTCgCGACGCACGGATCCGCTGTAGTTTTAGAGCTAGAAATAGCAAG | to construct pCFD4-sloth1 (aka JAB203) |
| JB573_CG32736gRNAdKO_R | ATTTTAACTTGCTATTTCTAGCTCTAAAACGGAAAGAAGCGCTTCGGTGTcGACGTTAAATTGAAAATAGGTC | to construct pCFD4-sloth1 (aka JAB203) |
| GP01169_F | TATATAGGAAAGATATCCGGGTGAACTTCGATGCCCGCCGGAGTTTCCTGGTTTTAGAGCTAGAAATAGCAAG | to construct pCFD4-sloth2 (aka GP01169) |
| GP01169_R | ATTTTAACTTGCTATTTCTAGCTCTAAAACAGGAATTTCAGGTACTGGCCGACGTTAAATTGAAAATAGGTC | to construct pCFD4-sloth2 (aka GP01169) |
| JB576_CG32736_CG42308gRNAdel1_F | TATATAGGAAAGATATCCGGGTGAACTTCGCGGTTATATAATTCCGTGTGTTTTAGAGCTAGAAATAGCAAG | to construct pCFD4-sloth1-sloth2 (aka JAB205, for dKO) |
| JB577_CG32736_CG42308gRNAdel1_R | ATTTTAACTTGCTATTTCTAGCTCTAAAACATAACTTGTCTAGTACACAGcGACGTTAAATTGAAAATAGGTC | to construct pCFD4-sloth1-sloth2 (aka JAB205, for dKO) |
| JB628_CG32-42_LH_EcoRI_F | cccttcgctgaagcaggtggGCCTCGTTGTTGGTGTAC | to amplify LHA |
| JB634_CG32-42_LH_Gal4SV40_R | gtagcttcatggtgTGTTGGTTTCGCCAATTTTAAG | to amplify LHA |
| JB635_CG32-42_Gal4SV40_LH_F | gaaaccaacacaccATGAAGCTACTGTCTTCTATC | to amplify Gal4-SV40 |
| JB636_CG32-42_Gal4SV40_loxP_R | acgaagttatAGACATGATAAGATACATTGATG | to amplify Gal4-SV40 |
| JB637_CG32-42_loxP_Gal4SV40_F | tatcatgtctATAACTTCGTATAATGTATGCTATAC | to amplify loxP-RFP-loxP |
| JB631_CG32-42_loxP_RH_R | gtcaccaataATAACTTCGTATAGCATACATTATACGAAGTTATACC | to amplify loxP-RFP-loxP |
| JB632_CG32-42_RH_loxP_F | acgaagttatTATTGGTGACTAAAGCTATTTAAGTG | to amplify RHA |
| JB633_CG32-42_RH_XhoI_R | actcgattgacggaagagccCTTCAGGGGATCAAGGAAC | to amplify RHA |
| JB265_Gibson_pEntr_1F | AAGGGTGGGCGCGCCGAC | to amplify pEntr backbone |
| JB266_Gibson_pEntr_1R | GGTGAAGGGGGCGGCCGC | to amplify pEntr backbone |
| JB517_CG32736_pEntr_F | ccgcggccgccccttcaccATGAGTCCGTACAGCGGATC | to construct pEntr_sloth1 |
| JB518_CG32736_pEntr_R | gggtcggcgcgcccacccttTTAGTTGGTGTTATTCGCGG | to construct pEntr_sloth1 |
| JB519_CG32736_nostop_pEntr_R | gggtcggcgcgcccacccttGTTGGTGTTATTCGCGGCT | to construct pEntr_sloth1 |
| JB404_CG42308_pEntr_F | ccgcggccgccccttcaccATGCCCGCCGGAGTTTCC | to construct pEntr_sloth2 |
| JB405_CG42308_pEntr_R | gggtcggcgcgcccacccttTTATGCAGATTCCGGTGGC | to construct pEntr_sloth2 |
| JB509_CG42308_nostop_pEntr_R | gggtcggcgcgcccacccttTGCAGATTCCGGTGGCTT | to construct pEntr_sloth2 |
| JB742_hSMIM4_gBlock (incorrect reverse seq) | ccgcggccgccccttcaccATGTTTACAAGGGCACAAGTTCGCCGGATACTGCAACGAGTACCAGGTAAACAGCGCTTTGGCATCTATCGCTTCCTGCCATTCTTTTTTGTACTCGGCGGTACTATGGAGTGGATAATGATTAAAGTTCGAGTGGGCCAGGAGACATTCTACGATGTCTATAGGCGAAAAGCTAGTGAACGCCAGTATCAAAGGCGATTGGAAGACGAGgggtcggcgcgcccaccctt | to construct pEntr_hSMIM4 |
| JB732_hSMIM4_pEntr_F | ccgcggccgccccttcaccATGTTTACAAGGGCACAAGTTC | to construct pEntr_hSMIM4 |
| JB733_hSMIM4_stop_pEntr_R | gggtcggcgcgcccacccttctattaCTCGTCTTCCAATCGCCTT | to construct pEntr_hSMIM4 |
| JB743_hSMIM4_nostop_pEntr_R | gggtcggcgcgcccacccttCTCGTCTTCCAATCGCCTT | to construct pEntr_hSMIM4 |
| JB526_hC12orf73_gBlock | CCGCGGCCGCCCCCTTcaccATGCCCGCGGGCGTGCCCATGTCCACCTACCTGAAAATGTTCGCAGCCAGTCTCCTGGCCATGTGCGCAGGGGCAGAAGTGGTGCACAGGTACTACCGACCGGACCTGACAATACCTGAAATTCCACCAAAGCGTGGAGAACTCAAAACGGAGCTTTTGGGACTGAAAGAAAGAAAACACAAACCTCAAGTTTCTCAACAGGAGGAACTTAAATAAAAGGGTGGGCGCGCCGACCC | to construct pEntr_hC12orf73 |
| JB548_entr_c12orf73_nostop_R | gcccacccttTTTAAGTTCCTCCTGTTGAG | to construct pEntr_hC12orf73 |
| JB549_entr_c12orf73_nostop_F | ggaacttaaaAAGGGTGGGCGCGCCGAC | to construct pEntr_hC12orf73 |
| JB725_CG3242_genomic_pEntr_F | ccgcggccgccccttcaccTCATTAGCGATGACAGCG | to construct pEntr_sloth1-sloth2 genomic |
| JB726_CG3242_genomic_pEntr_R | gggtcggcgcgcccacccttAAACGTGGCGTCTTTTGAATG | to construct pEntr_sloth1-sloth2 genomic |
| JB727_CG3242_transcript_pEntr_F | ccgcggccgccccttcaccAATCGAACAGCTGATTGCTG | to construct pEntr_sloth1-sloth2 transcript |
| JB728_CG3242_transcript_pEntr_R | gggtcggcgcgcccacccttCTTAAATAGCTTTAGTCACCAATAACTTG | to construct pEntr_sloth1-sloth2 transcript |
| JB761_pEntr_genomicCG32_CG42del_F | actaagcaaaCACTGTGTACTAGACAAGTTATTGGTG | to construct pEntr_sloth1-sloth2 genomic derivatives |
| JB762_pEntr_genomicCG32_CG42del_R | gtacacagtgTTTGCTTAGTTGGTGTTATTCG | to construct pEntr_sloth1-sloth2 genomic derivatives |
| JB763_pEntr_genomicCG32del_CG42_F | ctgaaaacacGCAAAATGCCCGCCGGAG | to construct pEntr_sloth1-sloth2 genomic derivatives |
| JB764_pEntr_genomicCG32del_CG42_R | ggcattttgcGTGTTTTCAGCCGGCGGTTATATAATTC | to construct pEntr_sloth1-sloth2 genomic derivatives |
| JB533_PD43265F | GAAAGAAGCGCTTCGGTGTC | qPCR primers for sloth1 |
| JB534_PD43265R | TCCACGTAGTTCTTCGCCTG | qPCR primers for sloth1 |
| JB540_PD43573F | AGGACTTGCGCGTCTACATC | qPCR primers for sloth2 |
| JB539_PD43573R | GATCCACCTGTGTGCTGTGT | qPCR primers for sloth2 |
| JB713_Rp49_F | ATCGGTTACGGATCGAACAA | qPCR primers for Rp49 |
| JB714_Rp49_R | GACAATCTCCTTGCGCTTCT | qPCR primers for Rp49 |
| JB717_Gapdh_F | CCAATGTCTCCGTTGTGGA | qPCR primers for Gapdh |
| JB718_Gapdh_R | TCGGTGTAGCCCAGGATT | qPCR primers for Gapdh |
| JB1110_CG32736_indel_1F | CCTTAAAATTGGCGAAACCA | PCR primers to genotype and sequence sloth1-KO fly lines and S2R+ cell lines |
| JB1111_CG32736_indel_1R | TAAAAAGAAGAGCGGCAGGA | PCR primers to genotype and sequence sloth1-KO fly lines and S2R+ cell lines |
| JB1114_CG42308_indel_1F | CGCGAATAACACCAACTAAGC | PCR primers to genotype and sequence sloth2-KO fly lines and S2R+ cell lines |
| JB1115_CG42308_indel_1R | ATGTAGACGCGCAAGTCCTC | PCR primers to genotype and sequence sloth2-KO fly lines and S2R+ cell lines |
| JB580_CG32736_CG42308_geno_1F | gagcagtcgccgaaatagtc | PCR primers to genotype and sequence dKO fly lines and S2R+ cell lines |
| JB587_CG32736_CG42308_geno_4R | tgaaaccctttccctgtcac | PCR primers to genotype and sequence dKO fly lines and S2R+ cell lines |
| JB787_CG3242_LHA_F | tcgaaaagttgtgcctgatg | PCR primers to genotype Gal4-KI flies (left homology region) |
| JB662_Gal4seq1R | agcggagaccttttggtttt | PCR primers to genotype Gal4-KI flies (left homology region) |
| JB659_3P3dsred_seq1F | ACTCCAAGCTGGACATCACC | PCR primers to genotype Gal4-KI flies (right homology region) |
| JB790_CG3242_RHA_R | cgatgagccggctataaaaa | PCR primers to genotype Gal4-KI flies (right homology region) |
