## Supplemental File 3 for "Two neuronal peptides encoded from a single transcript regulate mitochondrial complex III in *Drosophila*"

| Supplemental File 3 | | | | |
|---|---|---|---|---|
| **pEntr plasmid** | **Expression plasmid** | **Final plasmid name** | **Alternative name** | **Fly Insertion site** |
| pEntr_sloth1_stop | pWalium10-roe | pWalium10-sloth1 | UAS-sloth1 | attP2 |
| pEntr_sloth2_stop | pWalium10-roe | pWalium10-sloth2 | UAS-sloth2 | attP2 |
| pEntr_sloth2_stop | pValium10-roe | pValium10-sloth2 | UAS-sloth2 | attP40 |
| pEntr_hSMIM4_stop | pWalium10-roe | pWalium10-hSMIM4 | UAS-hSMIM4 | attP2 |
| pEntr_hC12orf73_stop | pWalium10-roe | pWalium10-hC12orf73 | UAS-hC12orf73 | attP2 |
| pEntr_sloth1-sloth2 transcript | pWalium10-roe | pWalium10-sloth1-sloth2 transcript | UAS-sloth1-sloth2 | attP2 |
| pEntr_sloth1-sloth2 genomic | pBID-G | pBID-{sloth1-sloth2} | {sloth1-sloth2} | attP40 |
| pEntr_Δsloth1-sloth2 genomic | pBID-G | pBID-{Δsloth1-sloth2} | {Δsloth1-sloth2} | attP40 |
| pEntr_sloth1-Δsloth2 genomic | pBID-G | pBID-{sloth1-Δsloth2} | {sloth1-Δsloth2} | attP40 |
| pEntr_BFP_nostop | pAWF | Act-BFP-FLAG | | |
| pEntr_sloth1_nostop | pAWF | Act-sloth1-FLAG | | |
| pEntr_sloth2_nostop | pAWF | Act-sloth2-FLAG | | |
| pEntr_BFP_nostop | pMK33-GW-SBP | pMK33-BFP-SBP | MT-BFP-SBP | |
| pEntr_sloth1_nostop | pMK33-GW-SBP | pMK33-sloth1-SBP | MT-sloth1-SBP | |
| pEntr_sloth2_nostop | pMK33-GW-SBP | pMK33-sloth2-SBP | MT-sloth2-SBP | |
| pEntr_BFP_nostop | pAWH | Act-BFP-HA | | |
| pEntr_sloth1_nostop | pAWH | Act-sloth1-HA | | |
| pEntr_sloth2_nostop | pAWH | Act-sloth2-HA | | |
| pEntr_RFeSP | pAWH | Act-RFeSP-HA | | |
| pEntr_CG10075 | pAWH | Act-CG10075-HA | | |
| pEntr_sloth1-sloth2 transcript | pMK33-GW | pMK33-sloth1-sloth2 transcript | MT-sloth1/2 | |
